# Genetic and Environmental Determinants of Streaming and Aggregation in *Myxococcus xanthus*

**DOI:** 10.1101/2024.11.06.622364

**Authors:** Trosporsha T Khan, Patrick Murphy, Jiangguo Zhang, Oleg A Igoshin, Roy D Welch

## Abstract

Under starvation conditions, a spot of a few million *Myxococcus xanthus* cells on agar will migrate inward to form aggregates that mature into dome-shaped fruiting bodies. This migration is thought to occur within structures called ‘streams,’ which are considered crucial for initiating aggregation. The prevailing traffic jam model hypothesizes that intersections of streams cause cell crowding and ‘jamming,’ thereby initiating the process of aggregate formation. However, this hypothesis has not been rigorously tested, in part due to the lack of a standardized, quantifiable definition of streams. To address this gap, we captured time-lapse movies and conducted fluorescent cell tracking experiments using wild-type and two motility-deficient mutant *M. xanthus* strains. By quantitatively defining streams and developing a novel stream detection mask, we show that streams are not essential for nascent aggregate formation, though they may accelerate the process. Moreover, our results indicate that streaming has a genetic component: disrupting only one of the two *M. xanthus* motility systems hinders stream formation. Together, these findings challenge the idea that stream intersections are required to drive aggregate formation and suggest that *M. xanthus* aggregation may be driven by mechanisms independent of streaming, highlighting the need for alternative models to fully explain aggregation dynamics.

## Introduction

Collective migration is a critical behavior observed across various organisms, including fish, birds, ants, and microbes, enabling them to survive, feed, and evade predators^1–7^. In the Gram-negative, soil bacterium *Myxococcus xanthus*, collective migration facilitates feeding in nutrient-rich environments and promotes survival during nutrient scarcity^8^. On solid agar surfaces under laboratory conditions, *M. xanthus* swarms exhibit this behavior by expanding or contracting in response to nutrient availability^9,10^. During vegetative growth, swarms expand outward across nutrient-rich media, while under starvation conditions, they contract inward, executing a developmental process culminating in the formation of aggregates that eventually mature into dome-shaped fruiting bodies. Each fruiting body is made up of approximately 100,000 cells^8,11,12^, of which a subset differentiates into spores, marking the organism’s final step in its adaptive response to starvation^11^.

During the aggregation phase of *M. xanthus* development, migrating rod-shaped cells often become locally aligned, moving in parallel to form patterns that resemble streams^13–17^. As these streaming regions expand within a swarm, they sometimes intersect. It has been hypothesized that these intersections play a crucial role in development^14,15^. This hypothesis is based on several nested assumptions: first, that cell movement slows or stops at these intersections due to crowding, similar to cars in a traffic jam^15,18^; second, that “jammed” cells signal their neighbors to stop moving via cell-contact signaling (C-signaling), causing further localized immobilization^19^; and third, that these clusters of stopped cells are the first stage of aggregate formation^18,20–22^. From these assumptions, a stepwise narrative emerges: ‘stream-dependent aggregation’ or the ‘traffic jam’ model (TJM)^14,15^ (Fig. 1): (1) cells align to form streams, (2) streams intersect, (3) intersections become jammed, (4) jammed cells signal neighbors to stop, and (5) stopped cells begin to aggregate. Although experimental data supports some aspects of this model, the causal relationships connecting each step remain speculative.

**Figure 1.**
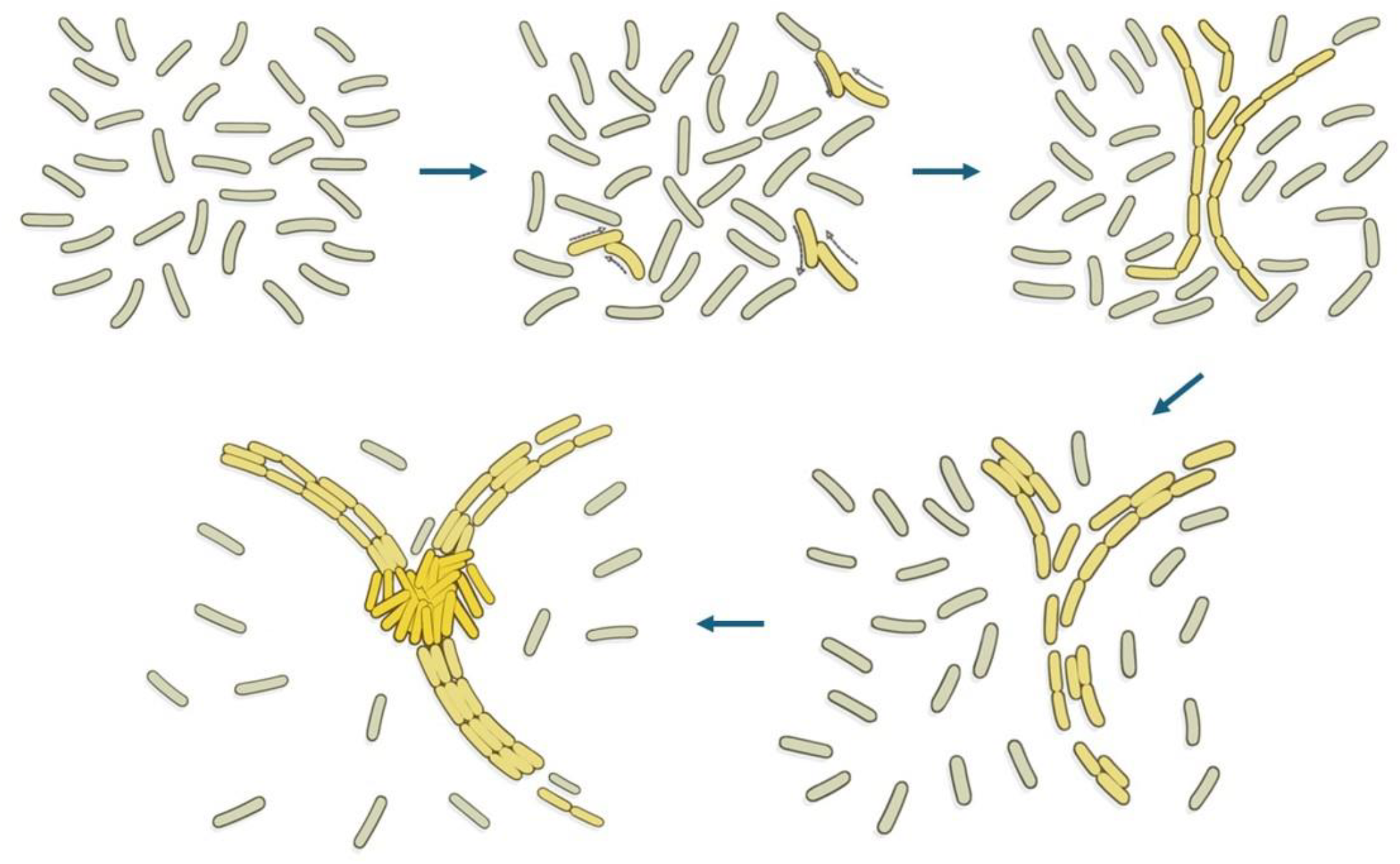
Stream-dependent aggregation model of *Myxococcus xanthus. M. xanthus* cells are described as self-propelled rods that collide with one another as they glide on the agar substrate. Collisions lead to cell alignment, forming long chains of cells in end-to-end contact. These chains align with each other forming streams. Streams are laid alongside other streams or at different angles, giving rise to stream intersections. Stream intersections are believed to be the points where traffic jams form, giving rise to nascent aggregates.

The most recent evidence supporting aggregate initiation through “traffic jam” comes from experiments that measure the physical properties of cells forming streams and initial aggregates. Streaming has been shown to be driven by mechanisms such as steric hindrance, random collisions, and slime trail following^23^. These mechanisms lead to streams characterized by nematically (i.e., in parallel) aligned cells moving together. At points of low alignment within these streams, known as topological defect points, half-integer topological charges (i.e., regions with asymmetric alignment contributing to polar clustering) can be observed^24^. Intrinsic cell polarity generates strong fluctuations in traction and cell flux, promoting the formation of layers around +1/2 defects^25^. While these data lend support to TJM, important conceptual gaps remain. Notably, defect points may not be the direct result of intersecting streams, and layers of cells within these defects may not in fact be nascent aggregates. If intersections are responsible for initiating aggregation, it would follow that all aggregates should be initiated at intersections.

An alternative model to stream-dependent aggregation involves chemotaxis-based aggregation, as observed in bacteria like *Escherichia coli*^26–28^ and the amoeba *Dictyostelium discoideum*^29,30^. In these organisms, chemotactic signals guide the initial positioning of aggregates and bias cell movement toward them. However, despite repeated attempts, no long-range chemotactic signal originating from *M. xanthus* aggregates has been identified. Although evidence exists for chemoattraction to phosphatidylethanolamine gradients^31^ in *M. xanthus*, this short-range morphogen has not been shown to bias *M. xanthus* streams toward aggregates.

Additionally, surface-associated signals such as contact-dependent C-signaling and exopolysaccharides (EPS) have been proposed to play a role in aggregation. End-to-end cell-contact via C-signaling may promote stream formation that coalesces into aggregates^32,33^. EPS, known to regulate cellular reversal and S-motility, is proposed to initiate aggregation through a positive feedback loop: collisions between cell groups increase local EPS concentration, reduce cell reversals, and trap cells within aggregates^34^. However, both models remain untested.

Occam’s razor might initially favor a chemotaxis-based aggregation model, yet the lack of genetic or biochemical evidence for an aggregate chemoattractant in *M. xanthus* has led to the adoption of the more complex stream-dependent aggregation model or TJM. The extent to which the TJM has become prominent is evident from a review of the scientific literature. A survey of published articles on *M. xanthus* development, motility systems, and aggregation mechanisms showed that about a quarter (31 out of 117) described aggregate initiation as cell “jamming” at stream intersections (Table S1). Thus, validating the TJM is crucial for refining current theories and ensuring that future models accurately reflect the biology.

Compounding the challenge of validating the stream-dependent TJM is the lack of a clear and quantifiable definition of streams in the current literature, which hampers our ability to identify, measure, or analyze streams or their intersections. Thutupalli et al (2015) demonstrated that cell reversal is necessary for stream formation^23^, but did not claim it was sufficient. Collective migration in *M. xanthus* is regulated by multiple genes^35–39^ associated with its two motility systems, Adventurous (A) and Social (S)^38,39^, as well as at least the *mgl*^40,41^ and *frz*^12,42^ gene clusters. However, except for the requirement of an intact *frzE* gene^23^, the role of other genes in stream formation remains ambiguous.

To begin validating the TJM in its entirety, its core assumptions must be examined. Streams must be quantitatively defined, their role in aggregate initiation must be clarified, and some of the genetic determinants must be identified and characterized. This study addresses these objectives by examining whether streams and stream intersections are necessary for aggregation and by assessing the contributions of *M. xanthus* motility systems to stream dynamics. By focusing on the initial phases of aggregation, when nascent aggregates first appear as regions of increased cell density, our approach enables us to investigate the dependence of aggregate initiation on stream formation and to determine whether stream intersections align with the timing and positioning of these nascent aggregates. Additionally, we assess the necessity of intact A- and S-motility systems for stream formation in *M. xanthus*.

## Results

### The minimal set of parameters required to define streams include cell movement, local cell alignment, local cell density, and stream persistence

Drawing from prior research that qualitatively describes *M. xanthus* streams, we aimed first to compile a ‘necessary and sufficient’ set of quantifiable parameters to characterize streaming regions within a swarm, ultimately identifying four: cell movement, local cell alignment, local cell density, and stream persistence^13,14,22,23,32,43–45^. To measure these four parameters, we captured time-lapse image stacks (1 frame/min) of wild-type *M. xanthus* DK1622 (WT) spotted at a relatively low cell density (LD) of 400 Klett (2.0 × 10^9^ cells/ml) from a nonhomogeneous cell suspension (i.e., some small clumps of cells remain after the final resuspension; see Fig. 2A, top left corner).

**Figure 2.**
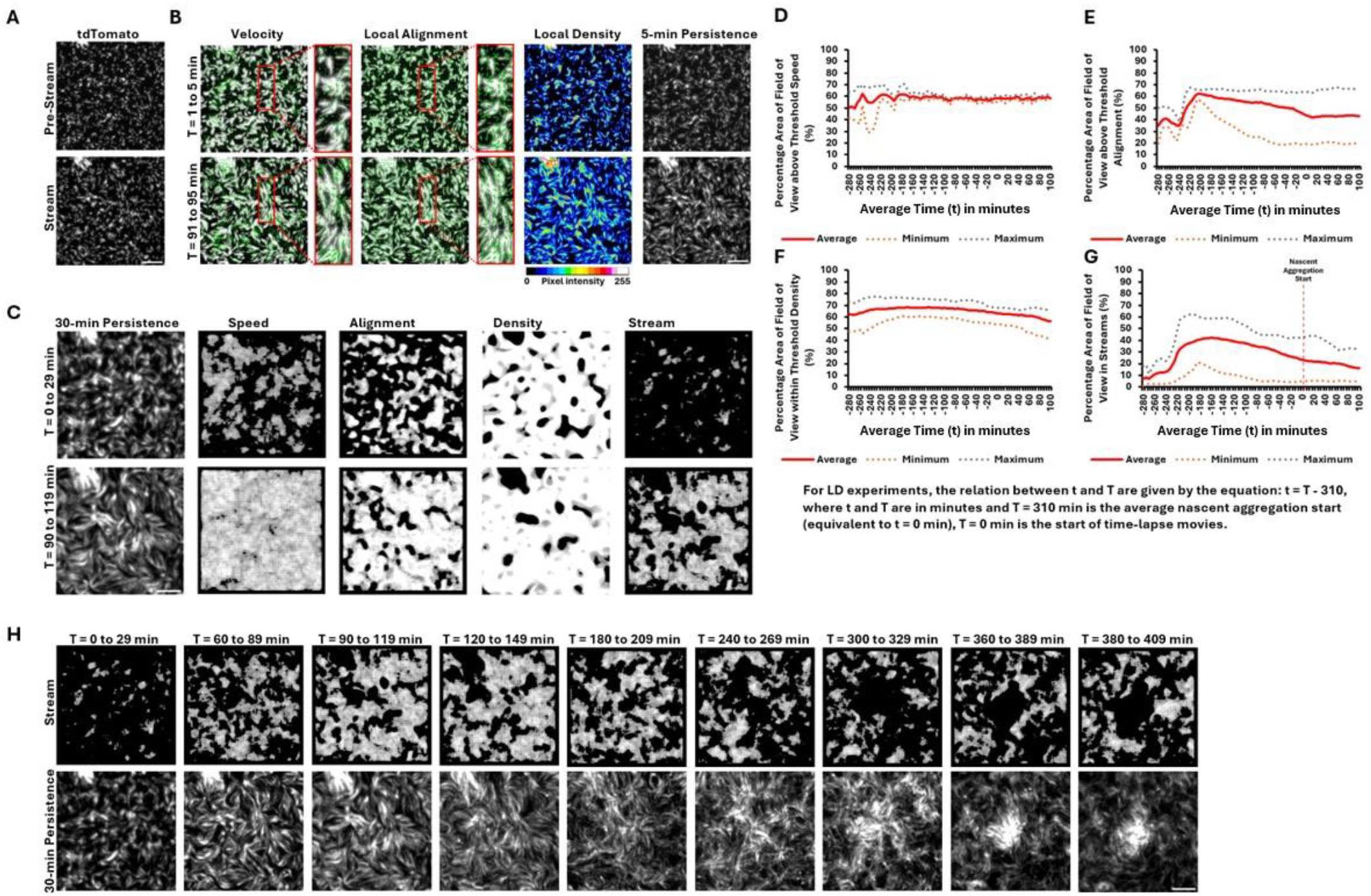
Stream mask generation to detect stream location in *M. xanthus*. (A) Visual representation of pre-stream and stream phase in LD time-lapse movies. A 400 by 400-pixel region of the original images are shown here. (B) Stream features used to generate stream masks. The stream phase compared with pre-stream phase indicates a greater number of cells with higher velocity (green velocity vector arrows), improved cell alignment (green alignment vector lines) along long axis of stream pathway, higher local density (pixel intensity) in streams compared to surroundings, and stream persistence for 5 minutes in the stream phase. The red boxes provide close-up view of cell velocity and alignment during the two phases compared. Time persistence images are derived from 1:20 dilution images, recorded in the tdTomato channel, by combining pixel intensities of five frames. Time persistence images are used to calculate velocity using Optical Flow method, alignment by Fourier Transformation, and density using Gaussian Filter. (C) A breakdown of the individual masks used to generate the stream mask. To generate individual masks, thresholds are applied to remove regions of low speed, low alignment, and lower- and higher-density regions. Black represents regions removed by the threshold. White represents regions meeting the required threshold. Stream masks capture regions where the thresholds for all four features, including time persistence, are met. During the stream phase a higher percentage of the field of view has attained threshold speed, local alignment, local density, and stream persistence, and therefore a higher percentage of streams. 30-min stream persistence is shown in the first column; to be qualified as a stream, the stream mask must detect its presence for at least 15 minutes. White represents streams and black indicates absence of streams. (D-G) Quantitative representation of the changes captured by individual masks over time. The data shown is an average, maximum and minimum of 50 samples from four replicate experiments. (H) Visual detection of stream formation over time using the stream mask. The highest stream detection is between 90 to 119 minutes in this example. Scale bar: 50 µm.

For these experiments, the initial timepoint (T = 0 min) corresponds to the first frame of each image stack, where cells had been on the agar surface for about five minutes, and they appeared randomly dispersed (Fig. 2A, top panel). During this initial ‘pre-stream’ phase, cells oscillated their leading pole with minimal total displacement (Movie M1). Over the next 30 to 90 minutes, cells transitioned into the ‘stream’ phase, where reversal frequency decreased, net cell movement increased and became more visually discernible, and cells began aligning into streams. The onset of the stream phase was also accompanied by an increase in local cell density (Fig. 2A, bottom panel).

Streams in WT *M. xanthus* appeared as regions where cells moved in parallel within high-density areas that persisted over time and space. Using the four parameters we identified—cell movement, local cell alignment, local cell density, and stream persistence—we defined and quantified streams in the observed images. These parameters were compared between the pre-stream (T = 1–5 min) and stream (T = 91–95 min) phases (Fig. 2B). As cells transitioned into the stream phase, velocity vectors increased in both magnitude and number, while alignment shifted from random to parallel lines along the axis of cell movement, resulting in nematic ordering similar to that of liquid crystals^23^. Streams also exhibited higher local density compared to their surroundings, as shown in the local density panel.

Finally, because streams are dynamic and transient, they may occupy a region for only a limited time. Thus, we measured their spatiotemporal persistence using cell tracks from the fluorescent channel (tdTomato), created by combining images over a period of five minutes. The five-minute persistence panels showed that in the pre-stream phase, both movement and alignment were low, whereas in the stream phase, cells exhibited high movement, alignment, and density (Fig. 2B). These persistence images contained measurable information on all four stream-defining features, making them suitable as input data to create a stream mask, which is a binary overlay that designates regions of interest in the image where stream formation and movement occur, distinguishing these areas from the surrounding background.

### Stream masks generated from stream defining parameters can detect stream location

The stream mask (Fig. 2C) identifies regions in the field of view (FOV) that meet or exceed thresholds for speed, alignment, and density, allowing us to visualize where streams exist. It also requires these features to persist in the same region for at least half of the input persistence time (e.g., 15 minutes in a 30-minute time-lapse). In the pre-stream phase, only small, disjointed regions met the criteria for streaming, whereas in the stream phase, continuous streams were detected, with low-speed, low-alignment regions and areas of extreme cell density (such as cell-free regions or initial clumps and aggregates) filtered out. The differences in mask outputs between these two phases support the validity of these parameters.

### The percentage of the FOV occupied by streams is closely influenced by the percentage of the FOV above threshold alignment in LD experiments

We applied the stream mask to time-lapse movies of LD to analyze changes in the percentage area of the FOV displaying stream properties over time (t). To distinguish between specific timepoints in the time-lapse and average timepoints in the graph, T and t are used respectively, where T = 0 min marks the start of the time-lapse and t = 0 min represents the average timepoint of nascent aggregation initiation. The graphs represent the percentage area of the FOV above threshold speed (Fig. 2D), alignment (Fig. 2E), and within threshold density (Fig. 2F) across time. A fourth graph illustrates the percentage area of the FOV classified as streams over time (Fig. 2G).

Initially, about half of the FOV has cells above threshold speed. This value fluctuates over the next 30 minutes, eventually rising to around 60% and then stabilizing (Fig. 2D). In the alignment graph, there is a substantial increase in the percentage area of the FOV above threshold alignment within the first 75 minutes, followed by a gradual decrease that levels off after nascent aggregation begins (Fig. 2E). The density graph shows a slight increase in the percentage area of the FOV within threshold density early on, followed by a slow decline as nascent aggregates form (Fig. 2F). The stream graph (Fig. 2G) reveals that the percentage area of the FOV in streams starts below 10%, rises to over 40% around t = -160 min (T = 150 min), then gradually decreases. By the time nascent aggregates appear, the percentage of the FOV in streams has halved to approximately 20%.

A notable observation is that the pattern in the stream graph closely resembles that in the alignment graph under the tested conditions (Fig. 2D–G). Since the speed mask primarily detects all cells in motion—including those moving at relatively low speeds—and the density mask captures all potential stream regions, including false positives, alignment and time persistence emerge as the key factors in distinguishing true stream regions. As a result, the shape of the alignment graph is strongly reflected in the stream graph.

### The beginning of the stream phase can be visually identified by applying the stream mask

The stream mask also allows us to pinpoint the timepoint of stream formation (Fig. 2H). Between T = 0 and 29 min, only small, disjointed spots are visible, indicating an absence of continuous streams. Streams begin to form between T = 60 and 89 min, becoming most prominent by T = 90–119 min. As time progresses, streams gradually decrease around T = 180– 209 min, while nascent aggregates become visible by T = 240–269 min. Maximum stream coverage occurs between T = 90 and 119 min, marking the transition from the pre-stream to the stream phase.

### Stream formation precedes aggregate initiation in WT

In WT LD experiments, stream formation consistently occurred before the initiation of aggregates (Fig. S1). To validate this observation, we compared the timepoint of maximum stream formation and the onset of nascent aggregation. In LD experiments, maximum stream formation was observed at t = -160 min (T = 150 min), whereas nascent aggregates began forming around T = 310 min (Fig. 2G, Table S2). This timing indicates that streams form in advance of aggregate initiation in LD experiment. New streams continued to form even after nascent aggregates had appeared.

Figure 3A visually illustrates stream formation preceding nascent aggregation. Streams are visible 30 minutes before nascent aggregate initiation, and when nascent aggregates appear, the corresponding regions are removed from the density and stream masks.

**Figure 3.**
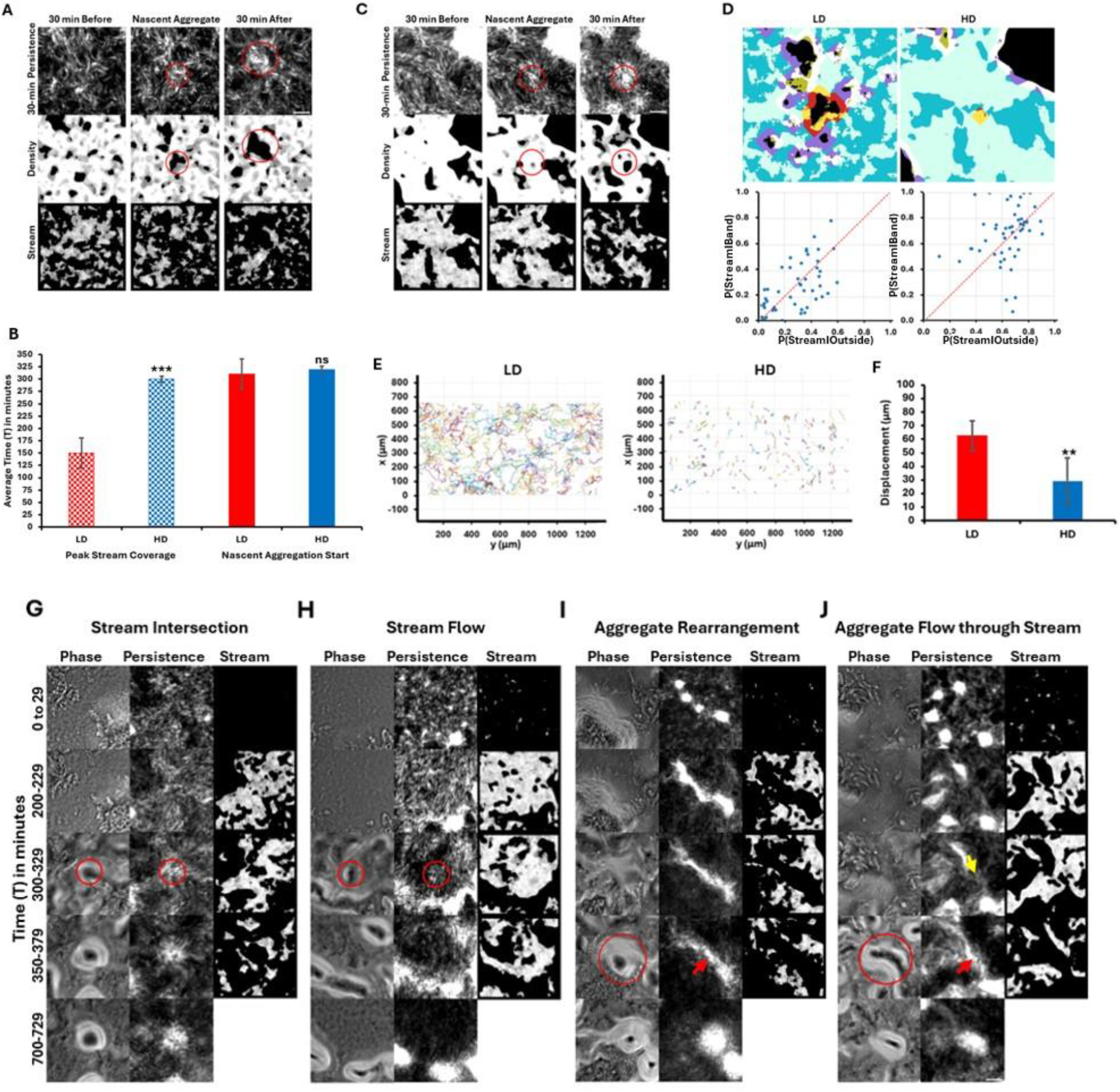
Nascent aggregate formation is not dependent on stream. (A-C) Visual representation of stream formation before nascent aggregate formation in LD and HD. (A and C) The first column shows 30 minutes before nascent aggregate formation, middle column shows nascent aggregate formation start, and the final column shows 30 minutes after. 30-min persistence, density mask and stream masks are shown. Red circles mark aggregates. (B) Comparison of timepoints of peak stream coverage and the time of nascent aggregate initiation. Average time (T) in minutes ± standard deviation from four replicate experiments is reported. Nascent aggregation starts at around the same time in LD and HD (P-value: 0.283), but LD shows peak stream coverage significantly earlier than HD (P-value: 0.001). (D) Proportion of streams around the aggregates compared to proportion of streams in the outside zone. All aggregates are marked black. The aggregate of interest is in the center with the region around it marked by a red band. The proportion of stream in this band (yellow) was calculated. All aggregates, whether developmental or non-developmental, were removed. The regions around the rest of the aggregates (marked purple) were also removed. The outside zone, devoid of aggregates and aggregate bands, was marked dark blue. The proportion of stream in the outside zone (light blue) was calculated. The proportion of stream in the band was plotted against proportion of stream in the outside for 50 aggregates of LD and HD experiments respectively. No significant difference was observed between the number of aggregates with higher P(streamIband) and higher P(streamIoutside). P-values: 0.85 for LD and 0.38 for HD, respectively. (E-F) Comparison of LD and HD cell tracks (E); and displacement before nascent aggregate formation (F). Displacement ± standard deviation from five replicate experiments is reported. HD shows a significantly lower displacement compared to LD. P-value: 0.01. (G-J) Visual representation of four representative mechanisms of nascent aggregation in HD. Red circles and arrows indicate nascent aggregates and yellow arrow shows aggregate direction of motion. Significance is determined by T-test and significance level is denoted by *p≤0.05, **p≤0.01,***p≤0.001 or ns for non-significance. Scale bar: 50 µm.

### Increased cell density does not accelerate nascent aggregation

To test whether increased cell density might accelerate nascent aggregate initiation, potentially causing aggregates to form before streams, we conducted high cell density (HD) experiments with WT at 2,500 Klett (1.25 × 10^10^ cells/mL), representing a 6.25-fold higher cell density than LD experiments (Movie M2).

Previous work by Liu et al. (2019) demonstrated that *M. xanthus* aggregation mechanisms shift with density, following a nucleation and growth pattern at lower densities and a spinodal decomposition pattern at higher densities^46^. We expected that such density-dependent shifts might also influence the timing and organization of stream formation and aggregation in HD conditions. At higher density, streams still formed before nascent aggregates (Fig. S2). The highest stream coverage (54% of the FOV), on average, was observed at t = -20 min (T = 300 min), while nascent aggregation began around T = 320 min (Fig. 3B, Fig. S3, Table S2).

Although there was only a 10-minute difference in nascent aggregate initiation times between LD and HD experiments, the timing of peak stream formation differed by 150 minutes (Fig. 3B). Comparison of all 50 HD nascent aggregation start times to the LD average (T = 310 min) shows that 80% of HD aggregates required equal or more time to initiate (Fig. S4). Figure 3C shows stream locations 30 minutes before, after, and at the time of nascent aggregation, confirming that streams precede aggregate initiation in both LD and HD conditions.

### Streams are not preferentially distributed around the nascent aggregates

We tested whether streams preferentially form around nascent aggregates by analyzing 50 nascent aggregates from both LD and HD experiments. The stream mask was applied 15 minutes before nascent aggregation, based on the hypothesis that streams guide cells toward aggregation centers^32^. We overlaid the stream and density masks (Fig. 3D, top panel) and compared the proportion of streams around the aggregate region (P(stream|band)) to those in the non-aggregate region (P(stream|outside)).

In both LD and HD conditions, nascent aggregates were nearly equally distributed on either side of the y = x line (Fig. 3D, bottom panel), indicating that the number of nascent aggregates with higher P(stream|band) is comparable to the number of aggregates with higher P(stream|outside).

A T-test confirmed no significant difference in the distribution of streams around nascent aggregates, suggesting that streams are randomly distributed rather than preferentially located around nascent aggregates.

### Cells do not rely on streams to reach nascent aggregates

Despite the random spatial distribution of streams, they consistently form before nascent aggregates in WT. To determine whether cells rely on streams to reach aggregation centers, we measured cell displacement prior to aggregate initiation. A common assumption regarding streams is that they facilitate long-distance cell travel toward aggregation points where cells then jam and aggregates initiate^10,22,23,33^. Thus, measuring cell displacement before nascent aggregation can indicate whether cells depend on streams to reach their aggregate location.

One-hour cell tracks beginning at T = 200 min are shown in Figure 3E. Under LD conditions, cells traveled relatively long distances, while in HD conditions, cells primarily moved back and forth over short distances. However, nascent aggregates formed at similar times in both conditions, suggesting that cells in HD do not require long-distance travel to initiate aggregation and therefore do not rely on streams for aggregation.

Given that *M. xanthus* cells reverse direction^47^, the actual displacement required for aggregate initiation is likely shorter than the total distance traveled. We estimated displacement in both LD and HD conditions from the beginning of the time-lapse until nascent aggregate initiation. On average, cells in LD moved 9 cell lengths (63 µm), while cells in HD moved only 4 cell lengths (29 µm) (Fig. 3F). The short displacement in HD suggests that cells from the immediate vicinity of nascent aggregates can directly contribute to aggregation, without the need for long-distance travel.

### Nascent aggregates form in multiple ways in HD DK1622 experiments, indicating that nascent aggregates and stream intersections are not always colocalized

In HD experiments, nascent aggregates initiated through multiple mechanisms, even in the presence of streams. We highlight four representative modes (Fig. 3G-J, Movies M3-M6), with additional modes described in Table S3.

Figures 3G and 3H depict stream-dependent aggregate initiation. In Figure 3G, an aggregate was initiated at the intersection of multiple streams, whereas in Figure 3H, one appeared within a single stream. The aggregate in Figure 3G persisted for at least 12 hours, while the one in Figure 3H eventually dissipated.

Figures 3I and 3J show aggregation initiated from other dissipating aggregates. In Figure 3I, cells redistributed among existing aggregates to form a larger diagonal aggregate, while in Figure 3J, cells extruded from dissipating aggregates to form connecting streams. One of these aggregates in Figure 3J relocated through a stream to initiate an aggregate in a new position. Despite the presence of streams, the primary contribution to nascent aggregates in these cases came from these preexisting ones.

### Streams are not required for aggregate initiation in A-S+ strain, and stream formation does not guarantee aggregate formation in A+S- strain

To further investigate the relationship between stream formation and aggregation, we examined two mutant strains: A-motility defective strain DK1218 (*cglB2-;* A-S+)^35^ that retains S-motility, and a S-motility defective strain DK1253 (*tgl1-;* A+S-)^36^, which retains A-motility^35,38^. In A-S+ strain (Movie M7), nascent aggregates formed without any streams, but they appeared much later in development (Fig. S5). Unlike WT, which forms streams prior to aggregation, A-S+ aggregates without forming any streams, suggesting that A-motility is essential for stream formation but not for aggregation. In contrast, A+S- strain (Movie M8), which lacks pili^36,38^ but retains A-motility, formed streams but did not form stable aggregates on TPM starvation agar (Fig. S5), highlighting that A-motility alone is insufficient to support stable aggregate formation.

We applied the stream mask to A-S+ before, during, and after aggregate initiation (Fig. 4A). No streams were detected, and aggregates only formed at later stages when compared to WT. When tested with the stream mask, A+S- showed weak streams initially (Fig. 4B), with the percentage of streams increasing in later frames but without stable aggregate formation, further differentiating its behavior from WT.

**Figure 4.**
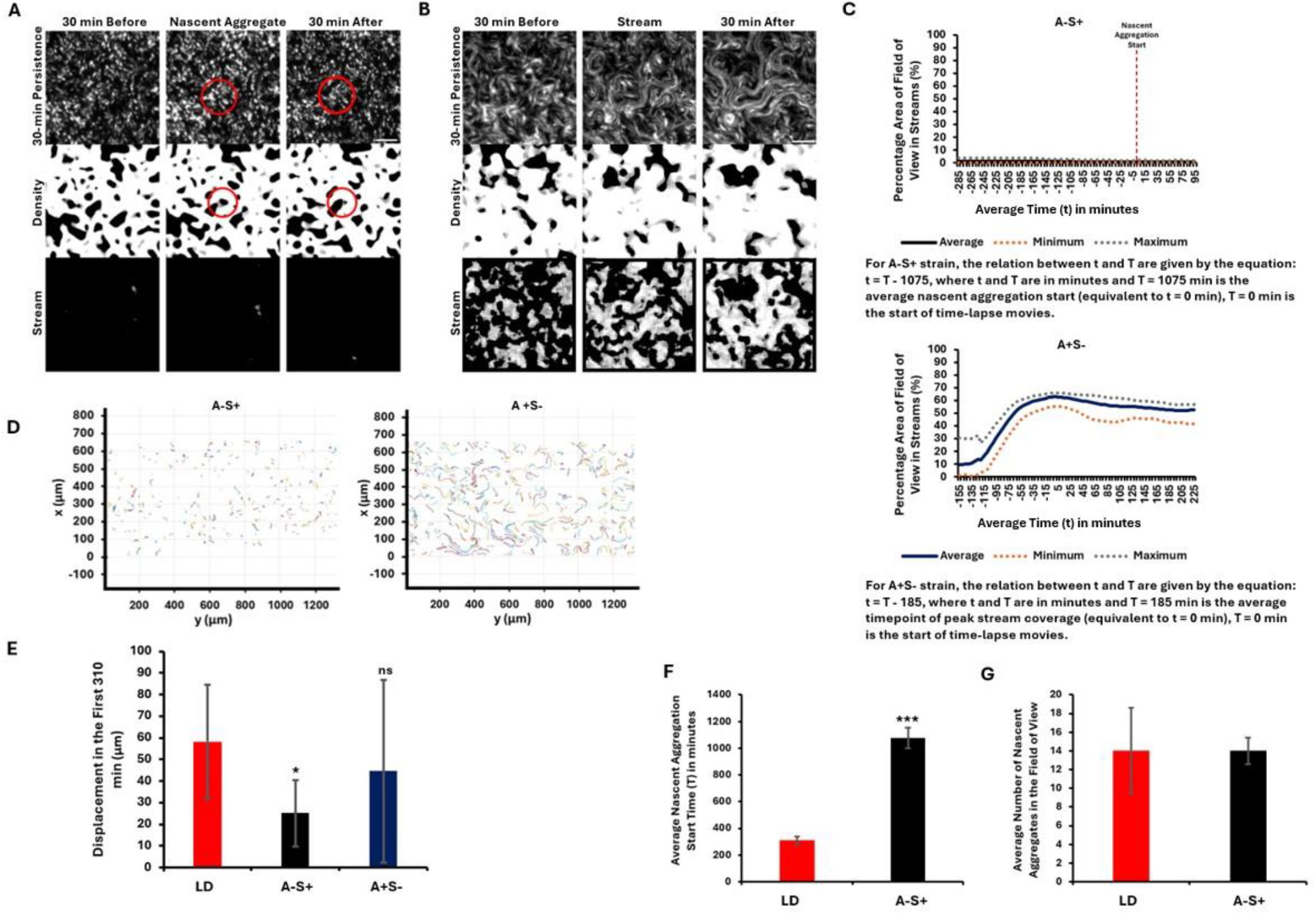
A-S+ forms aggregates without forming streams and A+S- forms streams but does not form aggregates. (A-B) Visual representation of nascent aggregate formation without stream formation in A-S+ (A) and only stream formation in A+S- (B). Middle columns show nascent aggregate initiation or stream, first columns show 30 minutes before, and final columns show 30 minutes after. 30-min persistence, density mask and stream masks are shown. Red circles mark aggregates. (C) Top panel shows percentage area of field of view in streams over time for A-S+, and bottom panel shows percentage area of field of view in streams over time for A+S- strain, calculated from the stream mask. The data presented is an average, maximum and minimum of 50 samples from four replicate experiments. (D) Comparison of A-S+ and A+S- cell tracks for 1 hour. (E) Displacement of LD compared with A-S+ and A+S- for the first 310 minutes of the cell-tracking movies. Displacement ± standard deviation from five replicate experiments is reported. T-test was performed. A-S+ shows a significantly lower displacement compared to LD. P-values: 0.04 and 0.29 for A-S+ and A+S-, respectively. (F-G) Comparison of nascent aggregation start time (F) and number of nascent aggregates (G) of LD and A-S+. Data represented is an average from four replicate experiments ± standard deviation. T-test was performed. A-S+ needs a significantly longer time to start developing nascent aggregates. P-value: 0.0002. Significance level is denoted by *p≤0.05, **p≤0.01, ***p≤0.001 or ns for non-significance. Scale bar: 50 µm.

Figure 4C compares the percentage of streams over time in the two mutants. A-S+ shows no streams. Nascent aggregates formed around T = 1075 min (Table S2). Quantification of alignment, density and speed masks are provided in Figure S6. A+S- showed an initial increase in the percentage of streams over time (Fig. 4C). The highest percentage was seen around T = 185 min (t = 0 min in the graph). After about 30 minutes, the percentage area of FOV in streams showed a small decrease which was also observed in the alignment and density mask quantifications (Fig. S6). It was observed that the streams in A+S-, after about T = 200 min, combined to form wider streams. The cells in these streams continued moving, forming tiering structures, but never formed stable aggregates as seen in WT (Movie M8, Fig S5).

### Stream-forming strains show a higher displacement compared to non-stream formers between the same timepoints

A comparison of cell tracking data between A-S+ and A+S- revealed that A+S- exhibited significantly higher movement compared to A-S+ for the same time period, starting from identical timepoints in the cell tracking movies (Fig. 4D). A+S- strain also exhibits greater heterogeneity in cell tracks compared to A-S+, LD, and HD (Fig. 3E), showing a mix of long and short tracks. In contrast, LD displays mostly long tracks, while A-S+ and HD show predominantly short tracks.

Cell displacements of A-S+ and A+S- were compared to wild-type DK1622 (LD) over the first 310 minutes (Fig. 4E). WT displayed the highest displacement (58 µm), followed by A+S- (45 µm) and then A-S+ (25 µm). Despite lacking S-motility, A+S- exhibited 1.8-fold higher displacement than A-S+. The high variability in A+S- tracks, noted earlier, is reflected in the large error bar, resulting in a relatively high average displacement and a non-significant difference from LD. Together, these comparisons highlight that, unlike WT, the mutant strains exhibit either stream formation without aggregation (A+S-) or aggregation without streaming (A-S+), underscoring the distinct genetic controls required for each process.

### Streams may improve the efficiency of aggregation by reducing the time needed for aggregate initiation

Finally, we investigated whether streams enhance the efficiency of nascent aggregation by shortening the time required for aggregate initiation. WT LD formed nascent aggregates faster than A-S+ strain, requiring 13 hours less time to reach initiation at the same cell density (Fig. 4F). Despite this time difference, both strains produced an equal number of nascent aggregates (Fig. 4G), suggesting that while streams might expedite aggregate initiation, they are not necessary.

## Discussion

In *M. xanthus*, development involves swarm self-organization into aggregates via a symmetry-breaking event^9^. Small behavioral or environmental variations may “seed” this process, which is amplified by communication and motility^32^. The traffic jam model (TJM), and subsequent models based on TJM, aligns with this view, proposing that streams precede aggregates and stream intersections act as symmetry-breaking seeds, impeding cell movement and initiating aggregation^13,14,22,24,32,48^. However, while stream intersections and aggregation sites may sometimes co-localize, this correlation does not prove causation. Our results do not support this expanded TJM.

Substantial research has focused on understanding the mechanisms underlying both stream^23,44,45^ and aggregate^15,17,18,33,46,49^ formation in *M. xanthus*. Thutupalli et al (2015) described the transition of cells from pre-stream to stream phases^23^ but did not establish a precise set of conditions necessary to define this transition. In our study, we aimed to determine the minimal parameters required to quantitatively define streams, enabling us to identify when and where they form. Using these parameters, we developed a novel stream detection mask, which we applied to various *M. xanthus* strains, successfully distinguishing between regions with and without streams.

We conducted experiments using both LD and HD conditions. While we observed streams and stream intersections in WT cells at both densities, neither the stream structure nor the intersection patterns matched those described by the TJM in Kuner and Kaiser (1982)^14^. Most notably, under both LD and HD conditions, alignment within streams prior to aggregate initiation was minimal, and aggregation initiated throughout the swarm rather than exclusively at intersections (Table S3). Defects resembling stream intersections—such as those described by Copenhagen et al. (2021)^24^—were occasionally observed in the same location just prior to the appearance of a nascent aggregate. However, similar defects were also observed without any subsequent aggregation, and the majority of aggregates formed at sites where no obvious defect was present. The TJM suggests a direct causal relationship between defects and nascent aggregation, but we observed no consistent spatiotemporal co-occurrence. Nor did we find any distinguishing features, such as differences in cell speed, between streams near aggregates and those elsewhere in the swarm that could help predict where an aggregate would initiate.

The TJM posits that streams act as ‘roads’^15^ for cells, facilitating rapid^32^, long-distance movement toward growing aggregates^10,22,23,33^. Extending this analogy, one might expect higher cell speeds and greater displacements on “better” roads (i.e., within more highly aligned streams). However, our findings do not support this prediction. Under LD conditions, streams contained cells with lower alignment but higher speeds and greater displacements prior to aggregate initiation, while HD cells exhibited higher alignment, lower speeds, and reduced displacements (Table S4, Fig. S7-S8, Fig. 3F). To identify streams before aggregate initiation using the stream mask, alignment or speed thresholds had to be set low enough that likely false-positive stream regions were sometimes detected at the beginning of the time-lapse. Thus, the analogy of cells using streams as directional ‘roads’ guiding them to nascent aggregates does not align with observed cell behaviors in either LD or HD conditions.

The idea of long-distance travel through streams is further challenged by our observation that cells exhibited only short displacements under both LD and HD conditions before nascent aggregates appeared. The field of view (FOV) spanned ∼1330 µm (∼190 cell lengths), yet average displacements were only 9 and 4 cell lengths (63 µm and 29 µm) for LD and HD, respectively—at least 21 times smaller than the FOV. This suggests that current in silico models^16,22^, which assume cells traverse the entire FOV and randomly collide to form streams prior to aggregation, may need adjustment to reflect these shorter displacements.

Existing in silico models of *M. xanthus* development often begin with streams formed by self-propelled rods that eventually aggregate at intersections through physical interactions^16^. Our WT experiments challenge this assumption, and mutant analysis further suggests that streaming and aggregation may be genetically separable phenotypes, with stream formation not necessarily preceding aggregation. A-S+ cells formed aggregates without streams, while A+S- formed streams without stable aggregates. This suggests that adventurous motility is essential for stream formation, whereas social motility is not. These data strongly indicate that the genetic controls governing streaming and aggregation differ, pointing toward correlation rather than causation between the two processes.

The advantage of stream formation is evident in our comparison of LD and A-S+ experiments. Although both strains produced an equal number of aggregates, A-S+ showed a significant delay in aggregation initiation. This suggests that streaming may enhance aggregation efficiency by enabling faster initiation, likely due to the greater cell displacement observed in LD for the same duration.

Kiskowski et al. (2004) demonstrated in simulations that aggregates can form from small, aligned patches, with streams emerging later to redistribute cells among aggregates^50^. This mechanism resembles the current TJM, except that in the Kiskowski model, aggregates preceded streams. Over time, the traffic jam narrative has accumulated assumptions to better fit available data, yet our study indicates that some of these assumptions do not hold under experimental scrutiny. We therefore conclude that while the TJM initially appeared to explain some observable events in *M. xanthus* development in the absence of a chemotactic signal, it now requires reexamination. Given recent research, it may also be time to reconsider the exclusion of chemotactic or other surface-associated signaling as a factor.

## Methods

### Strains and culture conditions

*Myxococcus xanthus* DK1622^51^ was used as the wild-type (WT). Motility mutants DK1253 (*tgl1-;* A+S-)^36^ and DK1218 (*cglB2-*; A-S+)^35^ were used to test their ability to make streams and aggregates under starvation^38^. DK1253 is deficient in S-motility with mutation in *tgl1* gene^36^ and DK1218 is deficient in A-motility with mutation in *cglB2* gene^35^. For fluorescent image series, tdTomato-expressing strain LS3908^52^, RW1253Td and RW1218Td were used. RW1253Td and RW1218Td were made by electroporation of plasmid pLJS145^52^ into A+S- and A-S+ strains respectively to generate tdTomato-expressing A+S- and A-S+ cells. Briefly, pLJS145 harbors the *tdTomato* gene under the control of an IPTG-inducible promoter derived from the vector pMR3487^52^. Once electroporated, a 1.38-kb DNA fragment upstream the IPTG-inducible promoter allows chromosomal integration into *M. xanthus* genome through homologous recombination^53^. All cells were grown overnight in CTTYE broth [1% Casein Peptone (Remel, San Diego, CA, USA), 0.5% Bacto Yeast Extract (BD Biosciences, Franklin Lakes, NJ, USA), 10 mM Tris (pH 8.0), 1 mM KH(H_2_)PO_4_, 8 mM MgSO_4_] at 32°C with vigorous shaking. tdTomato-expressing strains LS3908, RW1253Td and RW1218Td were supplemented with 10 µg/mL oxytetracycline for selection and 1 mM isopropylβ-D-1-thiogalactopyranoside (IPTG) for induction of tdTomato expression.

For development assays, cells were harvested mid-log phase, washed twice in TPM starvation buffer [10 mM Tris (pH7.6), 1 mM KH(H_2_)PO_4_, 8 mM MgSO_4_] and resuspended in TPM buffer to a cell concentration of 2 × 10^9^ cells/mL (400 Klett) for a low cell density experiment (LD). For a high cell density experiment (HD), 1.25 × 10^10^ cells/mL (2500 Klett) was used. Experiments with DK1622 were performed at both cell concentrations. All experiments with A+S- and A-S+ strains were performed at low cell density concentration. LS3908, RW1253Td and RW1218Td cells were diluted 1:1000 or 1:20 into DK1622, A+S- and A-S+ strains respectively. A droplet of 5 µL cells was spotted on agar slide complexes, as previously described^54^, containing 1% agarose-TPM medium supplemented with 1mM IPTG. 1:20 experiments were used for stream detection using the stream mask and 1:1000 experiments were used for cell-tracking experiments.

Cell resuspensions of DK1622 and A-S+ strain often contained initial preexisting cell clumps that were not removed before spotting. Therefore, developmental experiments were performed using nonhomogeneous cell suspensions.

### Time-lapse capture

Imaging was performed on a Nikon Eclipse E400 microscope with a pco.panda 4.2 sCMOS camera and NIS-Elements software. For stream mask and cell tracking experiments, LS3908, RW1253Td and RW1218Td samples were imaged with 400 ms exposure with a Sola LED light source at 75% intensity. Control of the fluorescent filter wheel and autofocus mechanism was managed with a MAC6000 system filter wheel controller and focus control module (Ludl Electronic Products, Ltd.). Images of DK1622 at LD and HD, and A+S- strain in the phase contrast and tdTomato channels were captured every 60 seconds for at least 12 hours. A-S+ strain images were captured in a similar manner, but because it develops much more slowly, they were recorded for 18 hours.

### Cell behavior data extraction

To quantify cell behavior, we performed the same procedures found in Cotter et al., 2017 and Zhang et al., 2020 at 1:1000 dilution to track fluorescently labeled tdTomato cells and classify both cellular transitions between non-persistent and persistent states and reversals in the persistent state^52,55^. Each trajectory segment with a start and end defined by a state transition or reversal, called a cell run, was then labeled, and the cell position, orientation, speed, and local alignment to other cells were recorded in a database. Cell tracking experiments were repeated five times.

### Image processing for stream mask experiment input

1:20 dilution experiments of DK1622 at LD and HD, A+S- and A-S+ were used for stream detection. Each experiment was repeated four times. 50 nascent aggregate zones were selected from the time-lapse movies of LD, HD and A-S+ strain. Nascent aggregate zone was identified visually from the tdTomato channel based on regions of higher pixel intensity, using the phase contrast channel as a guide. With the aggregate of interest positioned in the center, a 400 by 400-pixel region was cropped off from the original images. For each aggregate of LD and HD, the first 410 images were analyzed. For A-S+ strain, images closer to the end of the time-lapse were selected as development in A-S+ is slow. In this case the last 410 or 510 images were analyzed.

For A+S- strain, an equal region was cropped off the original images making sure that at least one stream was present in the field of view (FOV). Because streams were visible in the tdTomato images by 200 minutes in most cases, analysis of the first 410 images was sufficient. Similar to the aggregate region analysis, 50 stream regions were selected in total.

The cropped fluorescent images were further processed prior to any data extraction. First, stacks of images over a symmetric moving window of 5 frames in length were added pixel by pixel to create an image that captured persistence of cell movement. The python package skimage’s exposure method was used to perform histogram equalization on the resulting images to balance contrast in areas of low fluorescent intensity. Histogram equalized z-stack of 5 images were stacked into a larger stack of 30 consecutive images and computation was carried out at an interval of 5 minutes for the 410 or 510 minutes of the time-lapse as discussed above.

### Computational methods for stream mask generation

To estimate local velocity in the swarm of cells using optical flow, the approach of Vig, Hamby and Wolgemuth, 2016^56^ was applied to the images previously stacked and then normalized for contrast balance. Optical flow velocity of perceived movement in subregions of each image was calculated based on the changes in pixel intensity from frame to frame. While an orientation field can be obtained by normalizing the velocity vectors to unit length, this can create numerical issues when short vectors are scaled to have unit length. Any errors in the optical flow measurement for vectors with near-zero length will be magnified when the vectors are scaled up. Instead, the local alignment of the cells was estimated using 2D Fourier transforms as outlined below.

Local alignment was calculated using the code “Alignment by Fourier Transform” (https://github.com/OakesLab/AFT-Alignment_by_Fourier_Transform) by Marcotti et al., 2021^57^. The code works by first applying a fast Fourier transformation to the image in small windows, then using the frequency data to calculate the direction in which cells are aligned side-to-side in each window. The local nematic orientation of the biofilm is then the direction perpendicular to this calculated direction. Once the local orientation is known, the angles are used in a neighborhood to calculate an alignment order parameter in the range 0 to 1, with 0 being completely random alignment, and 1 being perfectly aligned. The code “AFT_batch.m” was run with a window of 15, a window overlap of 70%, and a neighborhood of 2 vectors to calculate the local alignment.

To isolate locations of streams, binary masks based on local density, local alignment, and velocity created were then combined. The mask for density was created first by using a Gaussian filter (imgaussfilt in MATLAB) with standard deviation of 10 pixels to blur the details of individual cells in the preprocessed images. The images then had regions of low or high fluorescence (e.g. mostly empty regions or aggregates, respectively) removed by applying lower and upper thresholds given in Table S4. Similarly, the masks for the speed and alignment were created by applying appropriate thresholds for each (Table S4), with the intention of removing areas with low speed or low alignment.

Output images from all four masks (speed, alignment, density and stream) were obtained separately for all 1:20 dilution experiments of LD, HD, A+S- and A-S+ experiments. White regions in the output images from the masks represent areas where cells meet the threshold requirements. Black regions indicate that the threshold is not met. Percentage area of the FOV (white regions) in these images was then measured using Fiji. Briefly, the images are segmented by thresholding and an ‘Analyze Particle’ tool in Fiji was used to measure the area of the resulting binary images.

### Parameter threshold optimization

Images for optimization were selected based on whether streams were already visible at that time point. In all cases the threshold was set to get maximum output rather than maximum accuracy. The speed mask was applied to 8 stream images from four replicate experiments starting from a speed threshold of 0.1 and ending at 0.7. The graphical and visual representation is shown in Figure S7. Speed threshold was set up to detect cell movement rather than to set a speed cut off. Therefore, the threshold was selected such that the speed mask outputs at least 80 % of the FOV which was usually lower than the cut off indicated by the optimization graphs.

An alignment mask was optimized to select cells in streams and leave out cells at regions of topological defects or low alignments. The alignment mask was applied to 8 stream images from four replicate experiments starting from alignment threshold of 0.1 and ending at 0.9. The graphical representation is shown in Figure S8. The threshold was set at the point right before the percentage area of cells shows a sharp decline.

The local cell density mask makes use of two threshold cut-offs (Fig. S9-S12). The first one (lower density threshold) separates all cells from the background and the second one (upper density threshold) removes regions of high density which can be either developmental aggregates or initial non-developmental cell clumps, making sure only cells in potential streams are captured. The two thresholds were optimized separately. In addition to that, each replicate experiment was separately optimized for density thresholds due to large variations in density within the movies (due to non-homogenous cell suspension) and between replicates. For lower density threshold, three stream images were selected from each replicate movie, whereas for upper density threshold, three aggregate images were selected. The images were analyzed through a range of thresholds suitable for the respective experiment. The cut-off values were selected by a combination of visual screening of the density mask output and the cut-off points indicated by the graphs. An average of the three values were used in the final experiments. All replicates of the same experiments were optimized similarly. A+S- strain was optimized for lower density threshold only (Fig. S11) and A-S+ strain was optimized for both density thresholds (Fig. S12) in the same manner as LD and HD experiments. A-S+ strain was tested for streams under the same speed and alignment threshold as LD.

### P(streamIband) and P(streamIoutside) calculation

For each aggregate, the initial formation time of the aggregates was identified. These aggregates were detected in fluorescent images using an intensity threshold. To study adjacent areas, the identified aggregates were dilated by 20 pixels, forming a ‘near-aggregate band’. Consequently, the images were divided into three distinct regions:

Aggregate (*S*_*aggregate*_): The area of the original aggregates.

Near Aggregate Band (*S*_*band*_): The area surrounding the aggregates.

Outside (*S*_*outside*_): All other areas of the image not included in the above regions.

Stream Region Identification: The region of streams *S*_*stream*_ was identified 15 minutes prior to the initial aggregate formation, using similar image processing techniques.

Proportion Calculations: To quantify the distribution of streams in relation to the aggregates, we calculated the proportions of streams as follows:

Proportion of Streams within the Band (P(stream|band)): This was calculated using the formula:

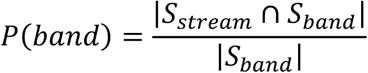

where |*S*_*stream*_ ∩ *S*_*band*_| represents the intersection of stream and band areas, and |*S*_*band*_| is the total area of the band.

Proportion of Streams outside the Band (P(stream|outside)): Calculated similarly, this is given by:

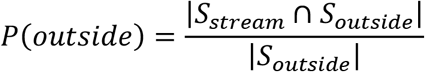

where |*S*_*stream*_ ∩ *S*_*outside*_| indicates the intersection of stream and outside areas, and |*S*_*outside*_| is the total area of the outside.

## Data Availability

The datasets generated and analyzed during the current study are available from the corresponding author on reasonable request.

## Acknowledgements

This research is funded by NSF MCB 1856665 and NSF DMS-NIGMS 1903160 (PI: RDW) and NSF DMS-1903275 and NSF IOS-1856742 (PI: OAI). The authors acknowledge Briquenne Williams for cartoon illustration of stream-dependent aggregation model, Niaz Zaidi for partial automation of alignment computation code and Laura Welch for helping in editing the manuscript.

## Author Contributions

Conceptualization and experimental design: RDW, TTK, PM; Data acquisition: TTK; Data analysis and interpretation: TTK, PM, JZ, OAI, RDW; Writing and revision: TTK, PM, JZ, OAI, RDW.

## Additional Information

**Supplementary information** accompanies this paper.

