## Supplementary Figures and Tables for "Genetic and Environmental Determinants of Streaming and Aggregation in *Myxococcus xanthus*"

**S1**

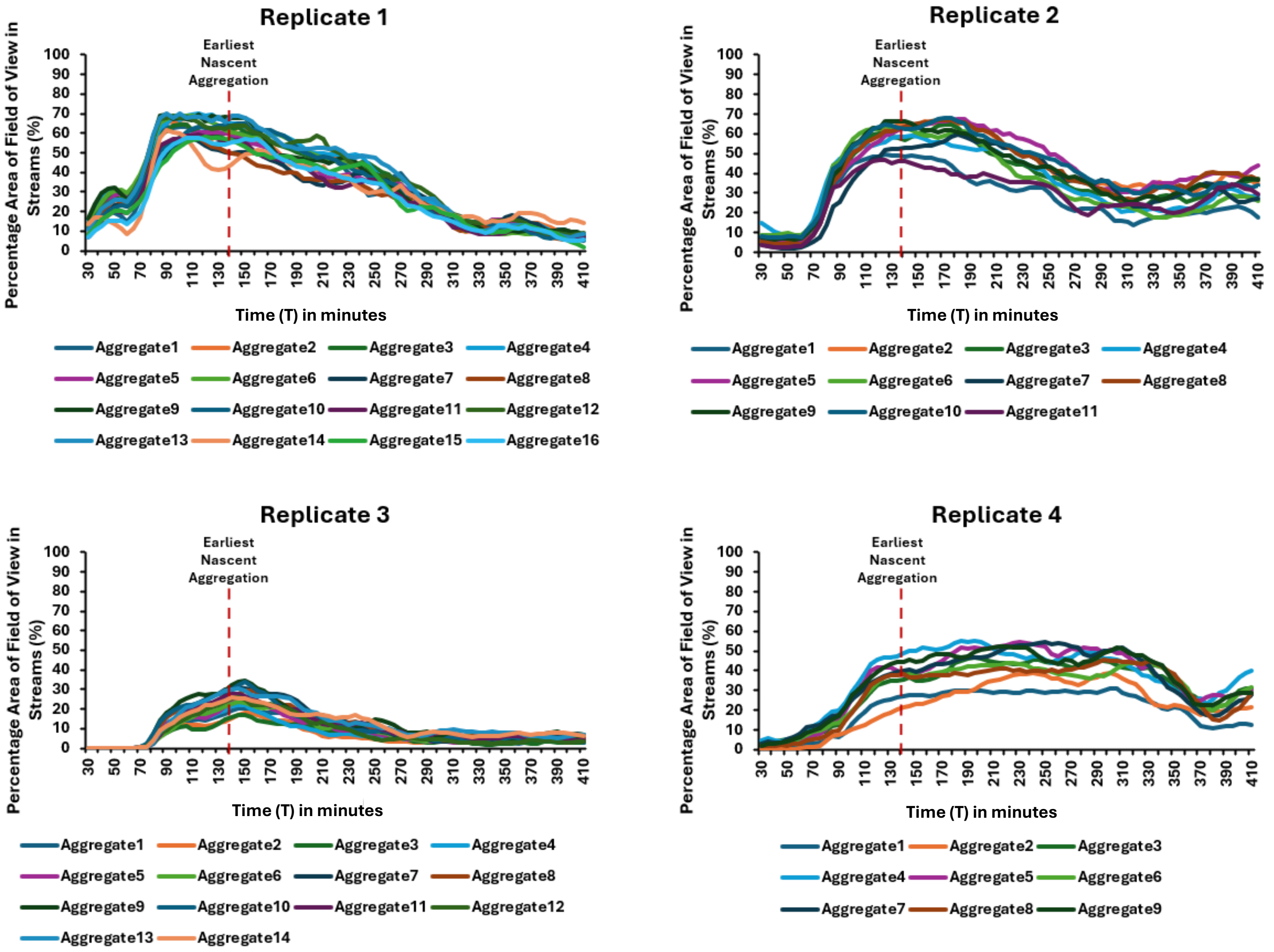


Fig S1. Streams formation precedes nascent aggregation in LD experiments for all 50 aggregates. Quantitative representation of changes in the percentage area of FOV in streams in LD experiment is shown from T = 30 min to T = 410 min for all 50 samples from four replicate time-lapse movies. The earliest nascent aggregation is observed at T = 140 min (broken red line) and the latest nascent aggregation is at T = 445 min. Note that in all cases, streams have formed before the earliest nascent aggregation time of T = 140 min. Here, T = 0 min is the first frame of the time-lapse movies.

**S2**

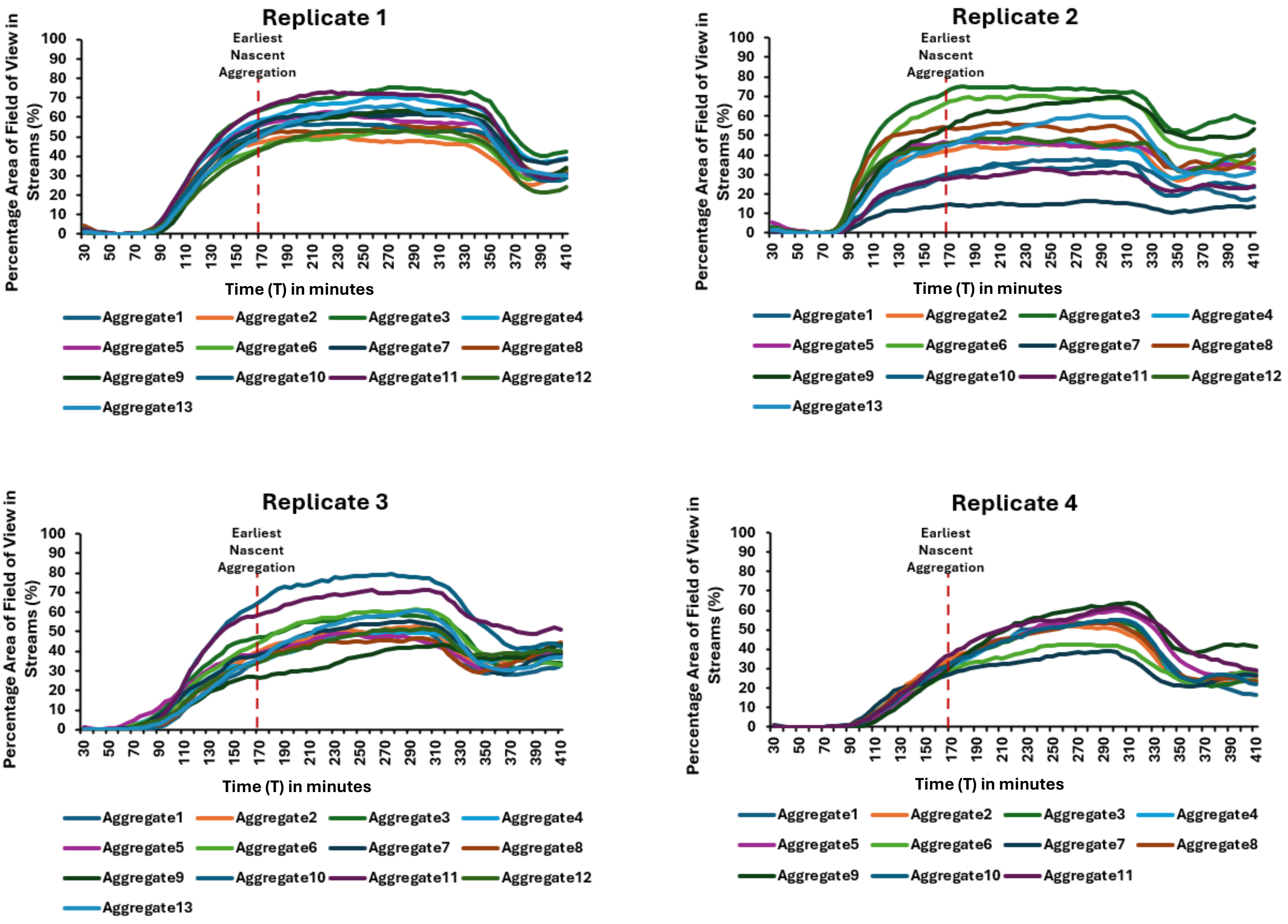


Fig S2. Streams formation precedes nascent aggregation in HD experiments for all 50 aggregates. Quantitative representation of changes in the percentage area of FOV in streams in HD experiment is shown from T = 30 min to T = 410 min for all 50 samples from four replicate time-lapse movies. The earliest nascent aggregation is observed at T = 170 min (broken red line) and the latest nascent aggregation is at T = 370 min. Note that in all cases, streams have formed before the earliest nascent aggregation time of T = 170 min. Here, T = 0 min is the first frame of the time-lapse movies.

**S3**

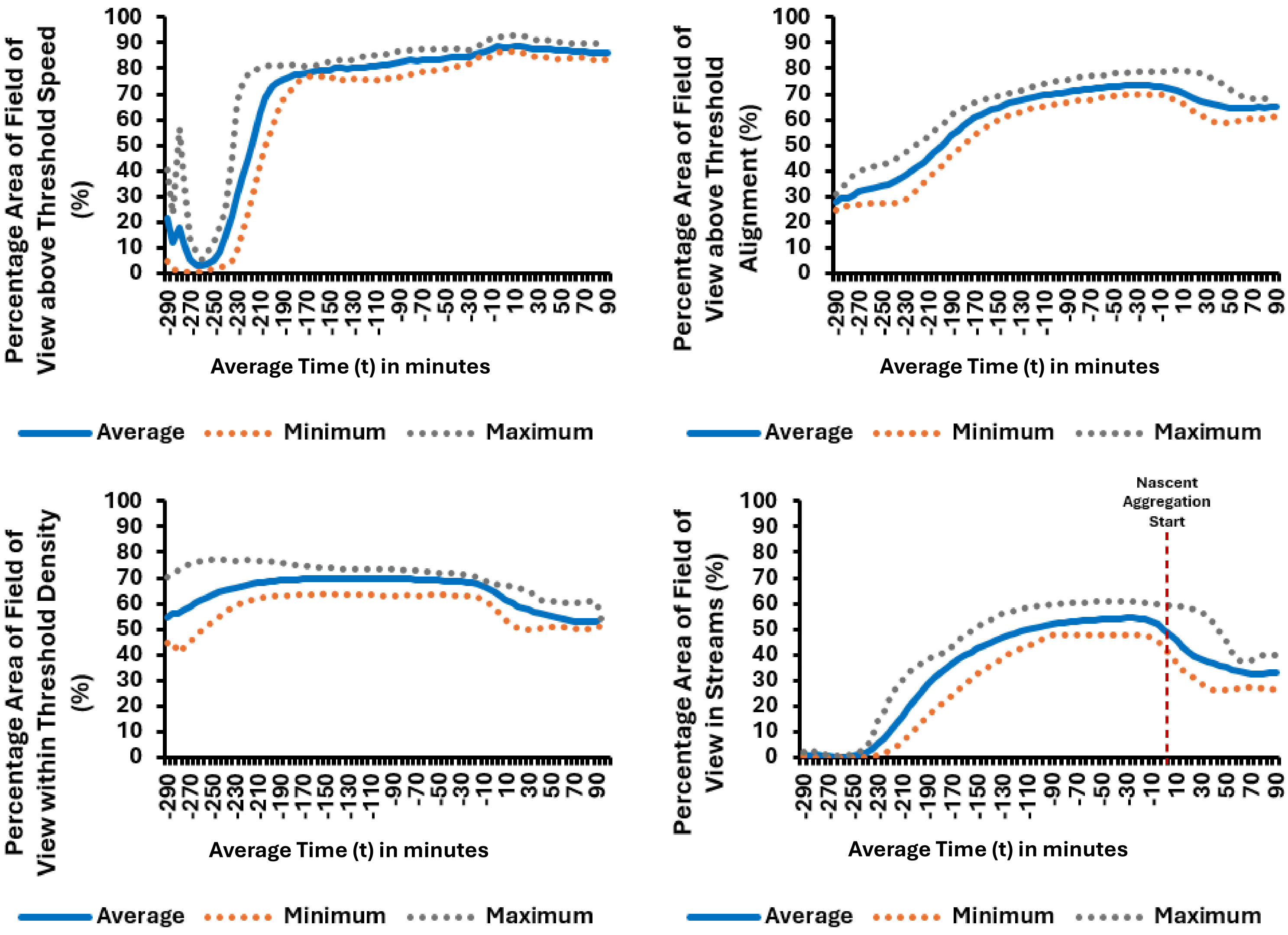


**For HD, the relation between t and T are given by the equation: t = T - 320, where t and T are in minutes and T = 320 min is the average nascent aggregation start (equivalent to t = 0 min), T = 0 min is the start of time-lapse movies.**

Fig S3. Quantitative representation of changes in the percentage area of FOV above threshold speed, local alignment, local density, and percentage area of FOV in streams over time in HD experiment. Nascent aggregation start is at t = 0 min in the graph or T = 320 min (red broken line), measured from the beginning of the time-lapse movies. The data shown is an average, maximum, and minimum of 50 samples from four replicate experiments.

**S4**

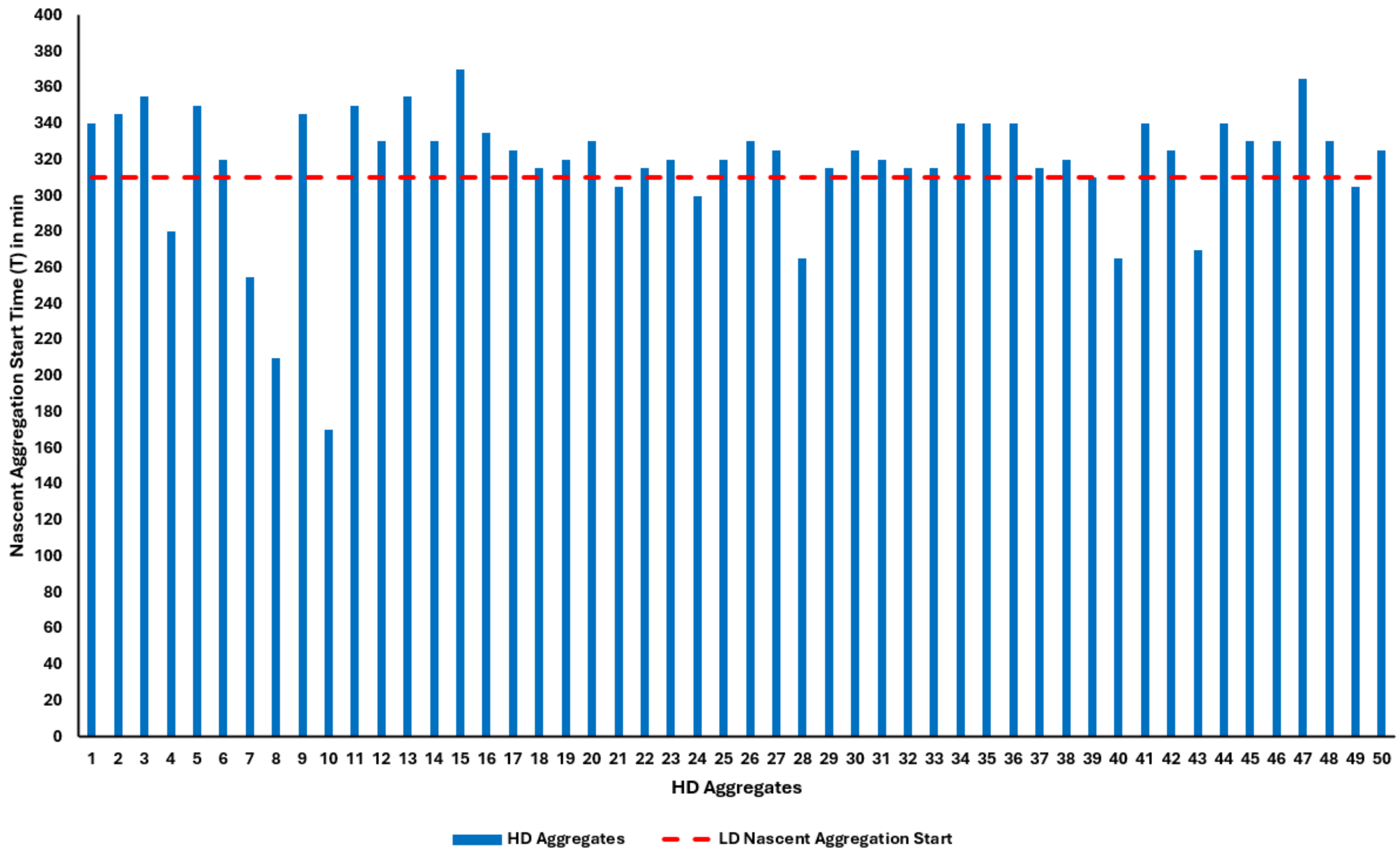


Fig S4. Nascent aggregation start time of all 50 HD aggregates from four replicate time-lapse movies are shown (blue bars). Broken red line indicates the average nascent aggregation start time in LD experiments. Out of the 50 HD aggregates, 40 aggregates (80%) initiated around the same time or later than the average nascent aggregation start of LD, indicating that increased cell density does not accelerate nascent aggregate initiation in HD.

S5

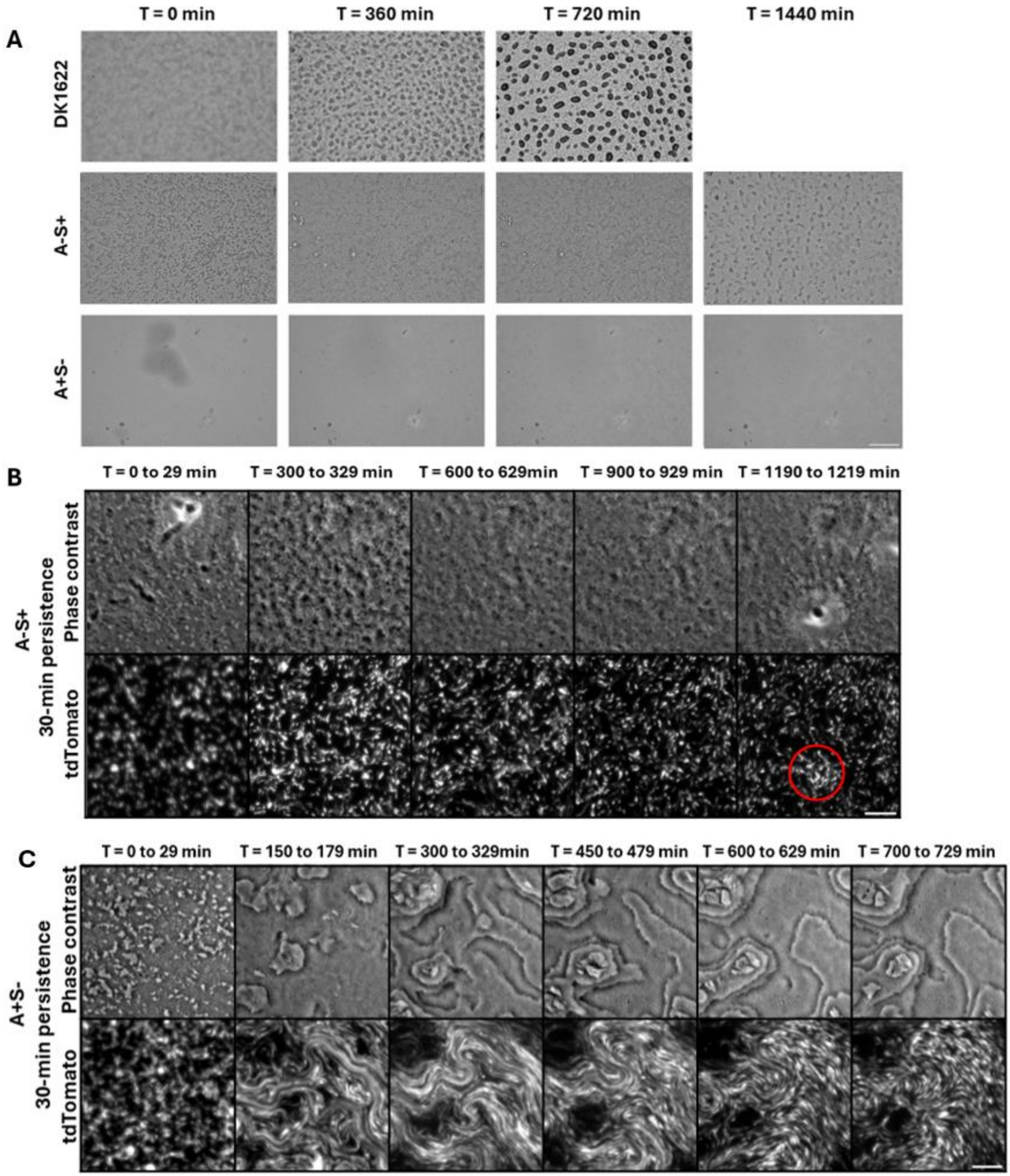
A
T = 0 min
T = 360 min
T = 720 min
T = 1440 min
DK1622
A-S+
A+S-
B
T = 0 to 29 min
T = 300 to 329 min
T = 600 to 629min
T = 900 to 929 min
T = 1190 to 1219 min
A-S+
30-min persistence
Phase contrast
tdTomato
C
T = 0 to 29 min
T = 150 to 179 min
T = 300 to 329min
T = 450 to 479 min
T = 600 to 629 min
T = 700 to 729 min
A+S-
30-min persistence
Phase contrast
tdTomato

Fig S5. (A) Bright field images of DK1622 compared with A-S+ and A+S-. DK1622 shows aggregates at both 360 and 720 minutes. DK1622 time-lapse was captured up to 720 minutes. A-S+ has a slower development and forms aggregates after 720 minutes in this example. Nascent aggregates are visible in the T = 1440 min frame (mature aggregates took longer to form and are not shown here). A-S+ time-lapse was extended up to 24 hours because of its slower development compared to DK1622. A+S- does not form aggregates or fruiting bodies. For A+S-, time-lapse was extended up to 24 hours to rule out the possibility of aggregation after 12 hours. Concentration of DK1622 and A+S- shown are at 3000 Klett and concentration of A-S+ shown is at 1000 Klett. Scale bar: 500 µm. (B) Time-lapse of A-S+ from phase contrast and fluorescent microscopy at 400 Klett. Nascent aggregate is seen at T = 1190-1219 min in the phase contrast. A red circle is drawn around the corresponding region of higher pixel intensity in the tdTomato channel. Scale bar: 50 µm. (C) Time-lapse of A+S- from phase contrast and fluorescent microscopy at 400 Klett. Phase contrast channel indicates tiering and tdTomato shows stream formation and no stable WT-like aggregation. Scale bar: 50 µm.

S6

A

B

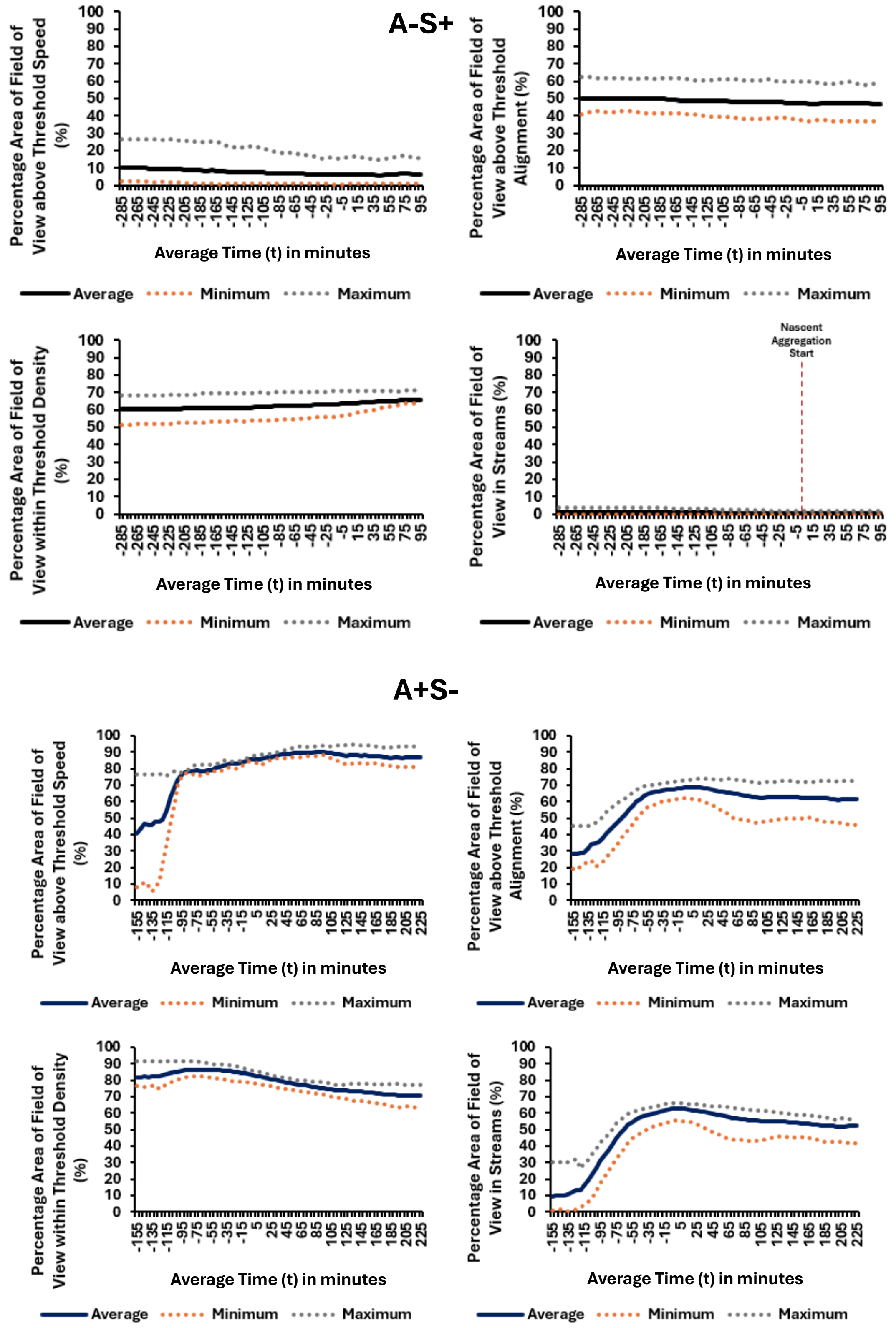


**For A-S+ strain, the relation between t and T are given by the equation: t = T - 1075, where t and T are in minutes and T = 1075 min is the average nascent aggregation start (equivalent to t = 0 min), T = 0 min is the start of time-lapse movies.**

**For A+S- strain, the relation between t and T are given by the equation: t = T - 185, where t and T are in minutes and T = 185 min is the average timepoint of peak stream coverage (equivalent to t = 0 min), T = 0 min is the start of time-lapse movies.**

Fig S6. Quantitative representation of changes in the percentage area of FOV above threshold speed, local alignment, local density, and percentage area of FOV in stream over time in A-S+ (A) and A+S- (B) experiments. The data shown is an average, maximum, and minimum of 50 samples from four replicate experiments of each type.

**S7**

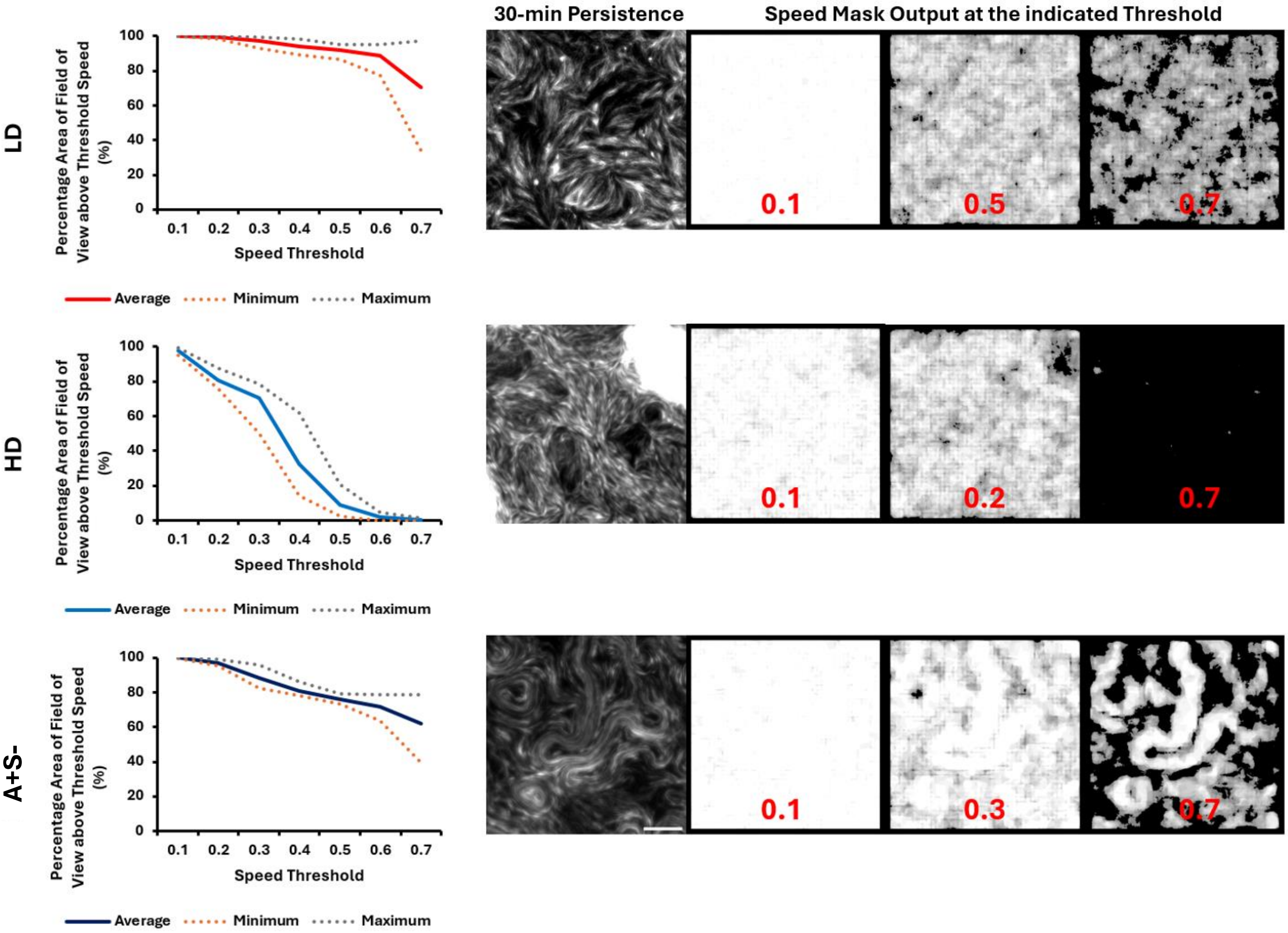


Fig S7. Speed threshold optimization for the speed mask in LD, HD, and A+S- experiments. The data presented is an average, maximum, and minimum of 8 samples from four replicate experiments of each type. Images are representatives of the time persistence and the output from the speed mask. 30 minutes persistence is shown for visual comparison. Speed mask output panel shows output at the lowest and highest threshold values (written in red over the images) tested with the output at the optimized value shown in the middle. Scale bar: 50 µm.

S8

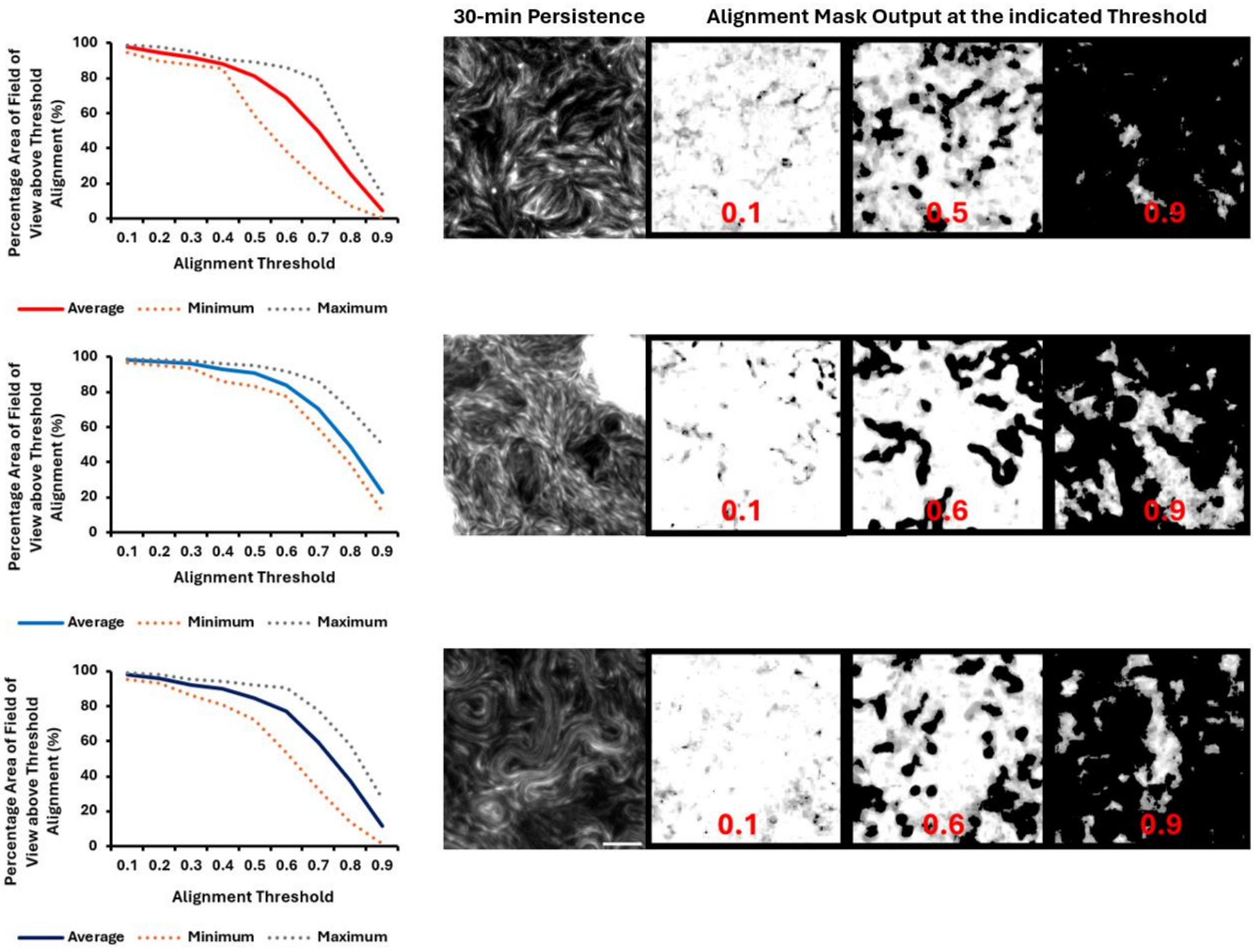


Fig S8. Alignment threshold optimization for the alignment mask in LD, HD, and A+S- experiments. The data presented is an average, maximum, and minimum of 8 samples from four replicate experiments of each type. Images are representatives of the time persistence and the output from the alignment mask. 30 minutes persistence is shown for visual comparison. Alignment mask output panel shows output at the lowest and highest threshold values (written in red over the images) tested with the output at the optimized value shown in the middle. Scale bar: 50 µm.

**S9**

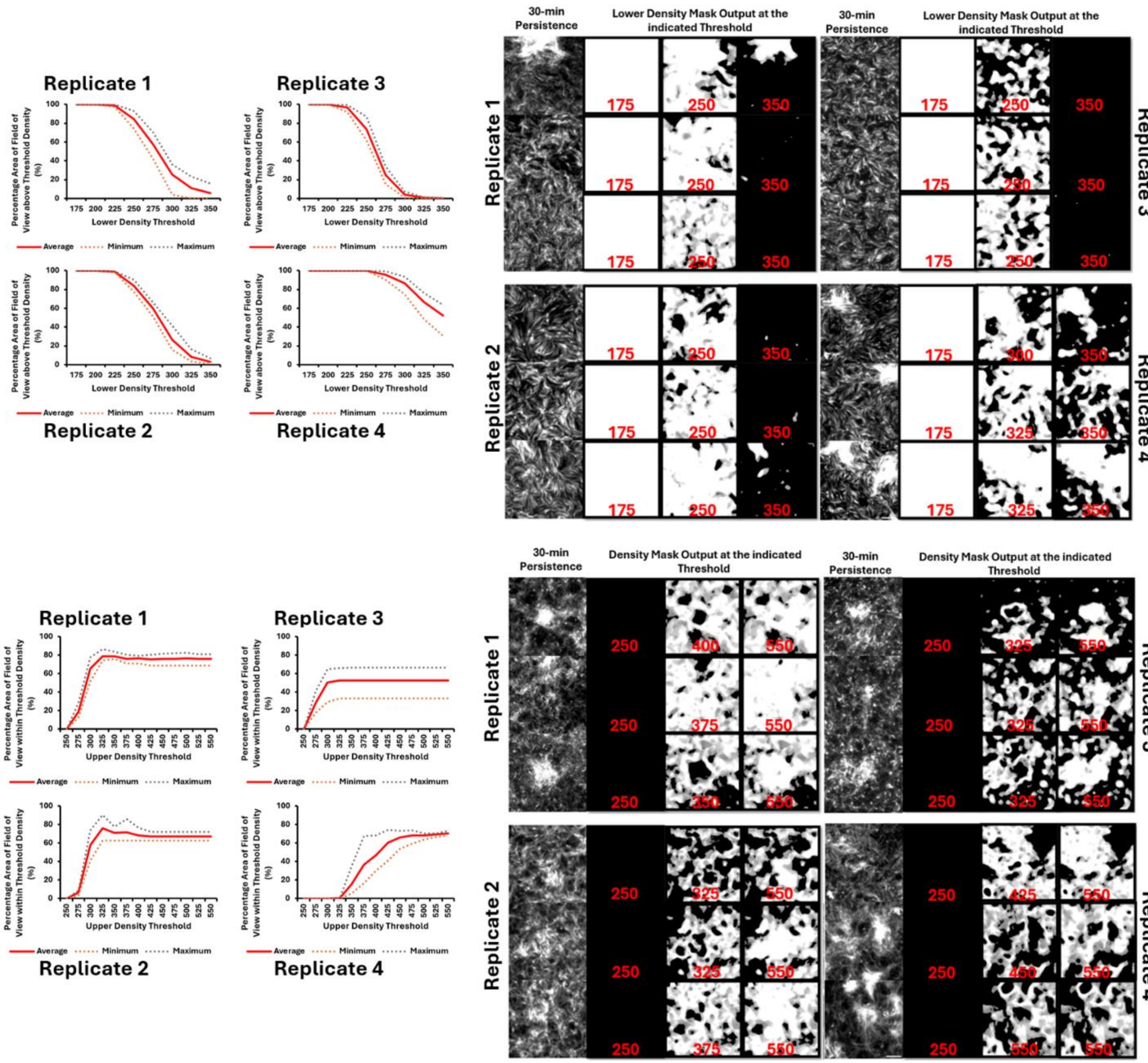


Fig S9. Density threshold optimization for LD experiments. Top panel represents lower density mask optimization and lower panel represents upper density threshold optimization. Three samples were chosen from each replicate experiment to perform optimization. An average of the three values obtained was used in the final experiment. Images are representatives of the time persistence and the output from the density mask. 30 minutes persistence is shown for visual comparison. Density mask output panel shows output at the lowest and highest values (written in red over the images) tested with the output at the optimized value in each case shown in the middle. Scale bar: 50 μm.

**S10**

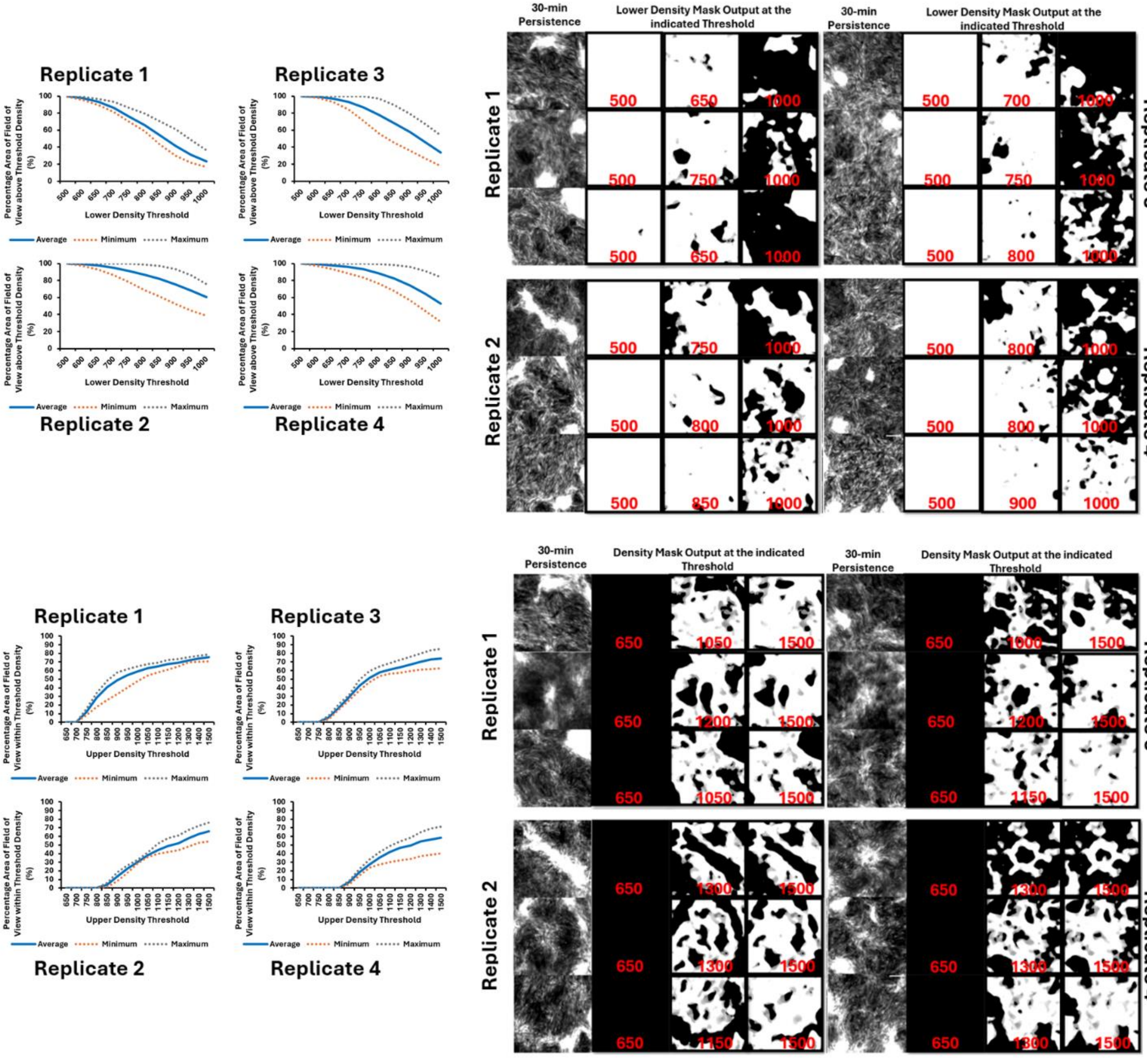


Fig S10. Density threshold optimization for HD experiments. Top panel represents lower density mask optimization and lower panel represents upper density threshold optimization. Three samples were chosen from each replicate experiment to perform optimization. An average of the three values obtained was used in the final experiment. Images are representatives of the time persistence and the output from the density mask. 30 minutes persistence is shown for visual comparison. Density mask output panel shows output at the lowest and highest values (written in red over the images) tested with the output at the optimized value in each case is shown in the middle. Scale bar: 50 μm.

**S11**

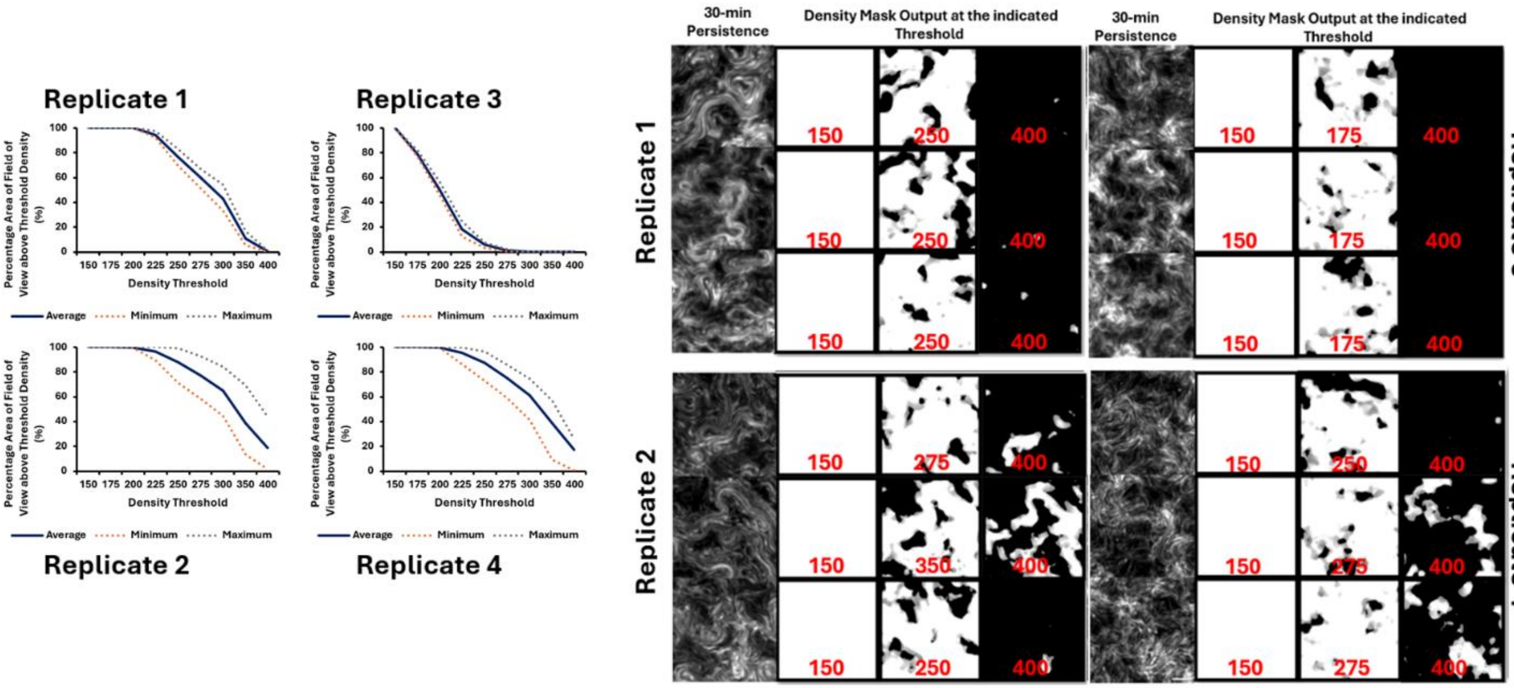


Fig S11. Density threshold optimization for A+S- experiments. Only a lower density mask optimization was performed since A+S- does not form stable WT-like aggregates. Three samples were chosen from each replicate experiment to perform optimization. An average of the three values obtained was used in the final experiment. Images are representatives of the time persistence and the output from the density mask. 30 minutes persistence is shown for visual comparison. Density mask output panel shows output at the lowest and highest values (written in red over the images) tested with the output at the optimized value in each case shown in the middle. Scale bar: 50 µm.

**S12**

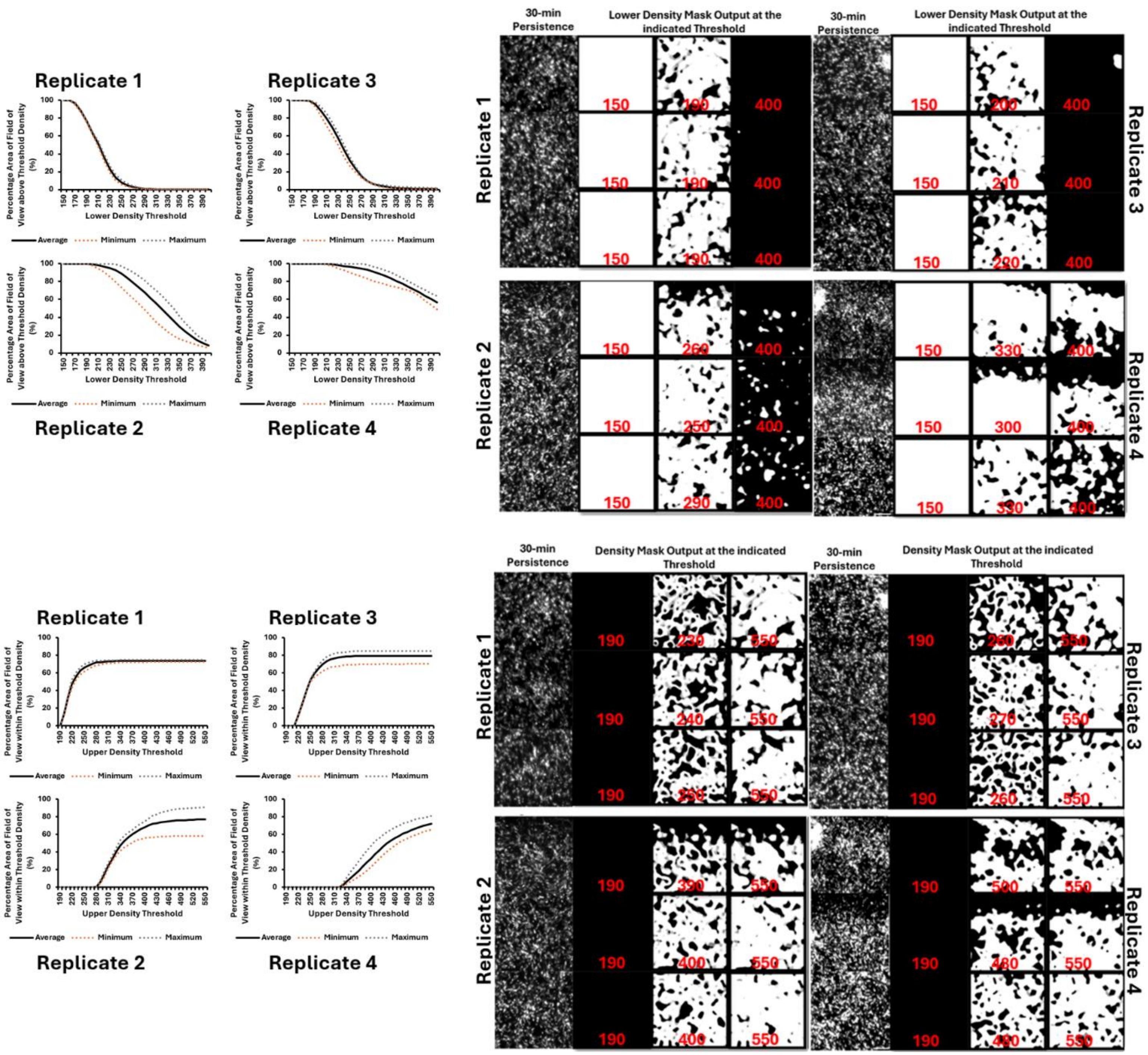


Fig S12. Density threshold optimization for A-S+ experiments. Top panel represents lower density mask optimization and lower panel represents upper density threshold optimization. Three samples were chosen from each replicate experiment to perform optimization. An average of the three values obtained was used in the final experiment. Images are representatives of the time persistence and the output from the density mask. 30 minutes persistence is shown for visual comparison. Density mask output panel shows output at the lowest and highest values (written in red over the images) tested with the output at the optimized value in each case shown in the middle. Scale bar: 50 µm.

**Table S1:** List of published articles linking aggregation to streams and stream intersections.

---

**Method:** We screened 117 published articles on *M. xanthus* searching for the words that have been used to describe streams or stream intersections. The search words used included- ‘stream’, ‘intersection’, ‘parallel’, 'align' ‘domain’, and ‘raft’. Any literature that described stream or stream intersection formation to precede aggregation or traffic jam formation was selected for this list, irrespective of the authors’ stance on the concept of aggregate initiation through traffic jam model. We found 31 papers that described aggregate initiation through traffic jam model in the introduction, results, or discussion sections. This list of 31 papers is provided below.

---

**Table S2:** Nascent aggregation start time in minutes for *M. xanthus* DK1622 at LD and HD conditions and for A-S+ strain.

| | Nascent Aggregation Start (min) | | |
|---|---|---|---|
| **Aggregate No.** | **LD** | **HD** | **A-S+** |
| **1** | 275 | 340 | 860 |
| **2** | 355 | 345 | 955 |
| **3** | 315 | 355 | 950 |
| **4** | 320 | 280 | 770 |
| **5** | 250 | 350 | 890 |
| **6** | 295 | 320 | 720 |
| **7** | 235 | 255 | 920 |
| **8** | 285 | 210 | 945 |
| **9** | 270 | 345 | 1110 |
| **10** | 340 | 170 | 1000 |
| **11** | 275 | 350 | 990 |
| **12** | 245 | 330 | 1200 |
| **13** | 310 | 355 | 1210 |
| **14** | 315 | 330 | 1055 |
| **15** | 370 | 370 | 1195 |
| **16** | 365 | 335 | 1050 |
| **17** | 355 | 325 | 1125 |
| **18** | 300 | 315 | 1050 |
| **19** | 305 | 320 | 930 |
| **20** | 445 | 330 | 1025 |
| **21** | 330 | 305 | 1025 |
| **22** | 265 | 315 | 990 |
| **23** | 270 | 320 | 1150 |
| **24** | 265 | 300 | 1130 |
| **25** | 260 | 320 | 1200 |
| **26** | 225 | 330 | 1015 |
| **27** | 330 | 325 | 1065 |
| **28** | 285 | 265 | 970 |
| **29** | 265 | 315 | 1220 |
| **30** | 360 | 325 | 1220 |
| **31** | 290 | 320 | 1145 |
| **32** | 445 | 315 | 1050 |
| **33** | 385 | 315 | 1220 |
| **34** | 395 | 340 | 980 |
| **35** | 340 | 340 | 1220 |
| **36** | 330 | 340 | 1220 |
| **37** | 360 | 315 | 985 |
| **38** | 375 | 320 | 1220 |
| **39** | 360 | 310 | 1040 |
| **40** | 350 | 265 | 1220 |

| | | | |
|---|---|---|---|
| **41** | 350 | 340 | 1220 |
| **42** | 140 | 325 | 1220 |
| **43** | 290 | 270 | 1220 |
| **44** | 315 | 340 | 1220 |
| **45** | 200 | 330 | 1090 |
| **46** | 340 | 330 | 1045 |
| **47** | 335 | 365 | 805 |
| **48** | 340 | 330 | 1140 |
| **49** | 355 | 305 | 1220 |
| **50** | 250 | 325 | 1180 |
| **Average** | 310 | 320 | 1075 |

**Table S3:** Multiple mechanisms of nascent aggregate initiation in *M. xanthus* DK1622 HD.

| HD Aggregate No. | Mechanism of Nascent Aggregation |
|---|---|
| **1** | Cells flow from aggregates in a stream-like fashion and from the surroundings in a non-stream fashion |
| **2** | Aggregate flow through streams |
| **3** | Cells flow from streams and the surroundings in a non-stream fashion |
| **4** | Cells flow from the nearest aggregate and stream flow |
| **5** | Cells flow from the nearest aggregates in stream-like fashion and surroundings |
| **6** | Cells flow from the nearest aggregates in stream-like fashion and from the surroundings in a non-stream fashion |
| **7** | Cells flow from all directions |
| **8** | Cells flow from aggregates, from surroundings and from streams |
| **9** | Cells flow from all directions and from an aggregate |
| **10** | Cells flow from aggregates, from surroundings and from streams |
| **11** | Cells flow from aggregates and from surroundings in a non-stream fashion |
| **12** | Cells flow from all directions including aggregates in both stream and non-stream fashion |
| **13** | Cells flow from all directions including aggregates |
| **14** | Aggregate rearrangement; initial aggregates join and cells from the surroundings also join |
| **15** | Cells from the streams and nearest aggregate |
| **16** | Cells flow from multiple directions including an aggregate |
| **17** | Stream flow |
| **18** | Aggregate formed from an initial aggregate and surrounding cells in stream and non-stream fashion |
| **19** | Stream flow side-by-side in opposite directions |
| **20** | Central aggregate rearranges; aggregate formed from an initial aggregate, surrounding cells, aggregates and streams |
| **21** | Aggregate flow through stream; cells flow mostly in opposite directions |
| **22** | Cells flow from aggregates and surroundings in a stream-like fashion |
| **23** | Cells flow in streams and non-streams |
| **24** | Aggregate flow through streams |
| **25** | Aggregate rearrangement; two aggregates combine and cells from surrounding join |
| **26** | Aggregates merge; streams and non-stream like flow |
| **27** | Cells move in a stream-like fashion |
| **28** | Possible stream intersection |
| **29** | Cells from the aggregates in a stream-like fashion |
| **30** | Cells in the initial aggregates rearrange by forming a stream intersection |
| **31** | Cells from aggregates; streams and non-streams |
| **32** | Cells from aggregates and streams |
| **33** | Stream intersection |
| **34** | Cells move in from all directions and streams |
| **35** | Cells from streams and surroundings |
| **36** | Stream intersection |
| **37** | Stream intersection |
| **38** | Stream intersection |
| **39** | Stream intersection |
| **40** | Stream intersection |
| **41** | Stream intersection and aggregate |
| **42** | Stream and non-stream movement |
| **43** | Stream intersection |

| | |
|---|---|
| **44** | Stream intersection |
| **45** | Cells from aggregates, mostly in a stream-like fashion; aggregate flow through stream |
| **46** | Cells from streams and non-streams |
| **47** | Stream intersection and surrounding cells |
| **48** | Aggregate cell rearrangement and cells from surrounding; streams and non-streams |
| **49** | Cells enter from all directions |
| **50** | Stream intersection |

**Table S4**: Parameter threshold optimization for *M. xanthus* DK1622 at LD and HD, A-S+ strain and A+S- strain,

| | Strain | | | |
|---|---|---|---|---|
| **Parameter** | **LD** | **HD** | **A-S+** | **A+S-** |
| **Speed threshold** | 0.5 | 0.2 | 0.5 | 0.3 |
| **Alignment threshold** | 0.5 | 0.6 | 0.5 | 0.6 |
| **Lower Density threshold, Replicate 1** | 250 | 685 | 190 | 250 |
| **Upper Density threshold, Replicate 1** | 375 | 1100 | 240 | -- |
| **Lower Density threshold, Replicate 2** | 250 | 800 | 275 | 290 |
| **Upper Density threshold, Replicate 2** | 345 | 1250 | 400 | -- |
| **Lower Density threshold, Replicate 3** | 250 | 750 | 210 | 175 |
| **Upper Density threshold, Replicate 3** | 325 | 1120 | 265 | -- |
| **Lower Density threshold, Replicate 4** | 320 | 835 | 320 | 270 |
| **Upper Density threshold, Replicate 4** | 475 | 1300 | 490 | -- |
