## Supplementary Video Legends for "Genetic and Environmental Determinants of Streaming and Aggregation in *Myxococcus xanthus*"

### Supplementary Videos

Supplementary Video 1: *M. xanthus* DK1622 (WT) development on TPM starvation agar at a low cell density (LD) of 2.0 x $10^9$ cells/mL. Randomly dispersed cells in the pre-stream phase oscillate, rearrange, and align parallelly to form streams. Peak stream coverage was observed between T = 95–124 min, and a nascent aggregate initiated around T = 260 min in this example.

Supplementary Video 2: *M. xanthus* DK1622 (WT) development on TPM starvation agar at a high cell density (HD) of 1.25 x $10^{10}$ cells/mL. Randomly dispersed cells in the pre-stream phase oscillate, rearrange, and align parallelly to form streams. Peak stream coverage was observed between T = 255–284 min, and a nascent aggregate initiated around T = 345 min in this example.

Supplementary Video 3: Example of a nascent aggregate initiation at the intersection of multiple streams under HD conditions. Peak stream coverage was observed between T = 260–289 min, and nascent aggregate initiated around T = 265 min.

Supplementary Video 4: Example of a nascent aggregate initiation within a single stream under HD conditions. This aggregate dissipated soon after its formation. Peak stream coverage was observed between T = 170–199 min, and nascent aggregate initiated around T = 315 min.

Supplementary Video 5: Example of a nascent aggregate initiating through cell redistribution from multiple preexisting non-developmental aggregates under HD conditions. Cells dissipating from these aggregates contributed to redistribution, giving rise to a nascent aggregate. Peak stream coverage was observed between T = 240–269 min, and the nascent aggregate initiated around T = 330 min.

Supplementary Video 6: Example of a nascent aggregate initiating through the relocation of a preexisting non-developmental aggregate via a stream under HD conditions. Cells dissipating from preexisting aggregates formed streams, one of which facilitated the relocation of an aggregate that gave rise to the nascent aggregate. Peak stream coverage was observed between T = 175–204 min, and the nascent aggregate initiated around T = 345 min.

Supplementary Video 7: *M. xanthus* DK1218 (A-S+) development on TPM starvation agar at a low cell density of 2.0 x $10^9$ cells/mL. Cells do not form streams, but a nascent aggregate was seen initiating around T = 1130 min (a timepoint much later than DK1622 at LD).

Supplementary Video 8: *M. xanthus* DK1253 (A+S-) on TPM starvation agar at a low cell density of 2.0 x $10^9$ cells/mL. Randomly dispersed cells in the pre-stream phase oscillate, rearrange, and align parallelly to form streams. Peak stream coverage was seen between T = 200–229 min. No nascent aggregate formed.
